# Confidence-based Prediction of Antibiotic Resistance at the Patient Level

**DOI:** 10.1101/2023.05.09.539832

**Authors:** Juan S. Inda-Díaz, Anna Johnning, Magnus Hessel, Anders Sjöberg, Anna Lokrantz, Lisa Helldal, Mats Jirstrand, Lennart Svensson, Erik Kristiansson

**Author notes:** Address correspondence to Juan S. Inda-Díaz, and Erik Kristiansson,.

## Abstract

Rapid and accurate diagnostics of bacterial infections are necessary for efficient treatment of antibiotic-resistant pathogens. Cultivation-based methods, such as antibiotic susceptibility testing (AST), are limited by bacterial growth rates and seldom yield results before treatment needs to start, increasing patient risk and contributing to antibiotic overprescription. Here, we present a deep-learning method that leverages patient data and available AST results to predict antibiotic susceptibilities that have not yet been measured. After training on three million AST results from 30 European countries, the method achieved an average accuracy of 93% across bacterial species and antibiotics. It predicted susceptibility with an average major error rate below 5% for quinolones, cephalosporins, and carbapenems, and below 8% and 14% for aminoglycosides and penicillins, respectively. Furthermore, the model predicted resistance with an average very major rate below 10% for cephalosporins, carbapenems, and aminoglycosides, but with higher very major error rates for penicillins and quinolones. We combined the method with conformal prediction and demonstrated accurate estimation of the predictive uncertainty at the patient level. Our results suggest that AI-based decision support may offer new means to meet the growing burden of antibiotic resistance.

**IMPORTANCE:** Improved diagnostic tools are vital for maintaining efficient treatment of antibiotic-resistant bacteria and for reducing antibiotic overconsumption. In our research, we introduce a new deep learning-based method capable of predicting untested antibiotic resistance phenotypes. The method uses transformers, a powerful AI technique that efficiently leverages both antibiotic susceptibility tests (AST) and patient data simultaneously. The model produces predictions that can be used as time- and cost-efficient alternatives to results from cultivation-based diagnostic assays. Significantly, our study highlights the potential of AI technologies to address the increasing prevalence of antibiotic-resistant bacterial infections.

## INTRODUCTION

The global rise of antibiotic-resistant bacterial infections threatens human health globally (*1*). In 2019, almost five million yearly deaths were attributed to antibiotic-resistant bacteria (*2*), a number that is expected to continue to grow in the coming decades (*3*). Reduced antibiotic efficacy in treatment increases the risk of performing vital healthcare procedures – including surgery, chemotherapy, and organ transplantation (*4*) – and, thereby, jeopardizes modern medicine as a whole.

Accurate and fast diagnostics are necessary for effective treatment of antibiotic-resistant bacteria. A central method is antibiotic susceptibility testing (AST), a laboratory test in which a bacterium isolated from a patient sample is cultivated and its resistance phenotype assessed in the presence of antibiotics (*5*). However, AST can be time-consuming due to the often low bacterial growth rate and the large number of antibiotics that may need to be tested for highly multidrug-resistant isolates. Tier-based testing strategies and time constraints usually yield incomplete AST results initially. For life-threatening infections, treatment needs to start as early as possible, often before AST result are available (*6*). Under these circumstances, the choice of treatment is reduced to educated guesses based on limited diagnostic information (*7*). This form of “empirical” treatment is associated with increased patient risks and overprescription of antibiotics (8–10).

Antibiotic resistance is commonly caused by resistance genes encoding various defense mechanisms. These genes are often co-localized on mobile genetic elements, such as plasmids and transposons (*11, 12*). Multiple resistance genes can thus be transferred simultaneously between bacterial cells, giving rise to strong correlations in susceptibility to different antibiotics. The infecting bacterium and, therefore, its susceptibility profile are, furthermore, dependent on patient characteristics, including age, sex, and the geographical region where the infection was acquired (13–15). Indeed, patient data has previously been shown to contain valuable information for selecting suitable antibiotic therapy for bacterial urinary tract infections (15–17). There are, however, no AI-based methods that can combine patient data with the initial, often incomplete AST results and thus guide treatment choice based on all available diagnostic information. Indeed, integrating patient data with available AST results could enable more accurate prediction of susceptibility to antibiotics that have not been tested, thereby providing physicians with comprehensive diagnostic information at an earlier point in time.

Artificial intelligence (AI) and deep learning have been successfully applied to diagnostics (*18, 19*), but the focus has primarily been on image-based data commonly used in radiology and pathology (*20*). In contrast, methods for non-image multimodal data, which are more prevalent in the diagnosis of infectious diseases, have received less attention (*21*). Several methods for predicting phenotypes from genotypic data have also been proposed (22), as have machine-learning methods using electronic health records and patient data (23). There are, however, few AI-based decision support systems for antibiotic treatment selection in use in clinical settings (*24*). A major culprit in the development of such methods is the complexity of the diagnostic data, which is typically categorical (stratified test results and patient data) and contains dependencies and redundancies. Also, when applied to antibiotic-resistant bacteria, the incompleteness of the diagnostic data, particularly AST results, requires approaches that can efficiently handle missing observations. Furthermore, since model accuracy depends on the amount of available information, any prediction must be accompanied by estimates of its uncertainty. Indeed, the possibility of disregarding predictions that are insufficiently confident is vital for critical decision-making and, thus, essential for the adoption of AI-based methodologies in healthcare settings (*25*). However, today, most AI-based diagnostic methods are primarily evaluated in populations and do not provide uncertainty information for patient-level predictions (*26*). Existing methods have, so far, been unable to adequately address these challenges and are either limited to a few specific antibiotics, cannot incorporate missing AST data, or do not provide any estimates of the predictive uncertainty (*27, 28*).

In recent years, transformer-based models, such as BERT (bidirectional encoder representations from transformers) (*29*) and GPT (generative pre-trained transformer) (*30*), have transformed natural language processing. These models operate on categorical input data, often structured into sentences of words, and subject them to multi-head self-attention (*31*). This process enables inference of complex word dependencies directly from data and thereby predicts missing parts. Therefore, we hypothesized that transformers may be suitable for predicting antibiotic susceptibility results from a combination of incomplete diagnostic information and patient data. Multimodal self-attention would enable the identification of complex dependencies among diagnostic data types and facilitate extrapolation to susceptibilities that have not been tested. Transformers have previously been shown to be highly useful beyond natural language processing (*32, 33*), but are rarely used for diagnosing infectious diseases.

In this study, we present a novel transformer-based method that accurately predicts antibiotic susceptibilities using patient data and incomplete diagnostic information for three common bacterial pathogens, *Escherichia coli*, *Klebsiella pneumoniae*, and *Pseudomonas aeruginosa*. We combine the model with conditional inductive conformal prediction (*34*) to estimate the uncertainty of each prediction at the patient level, thereby enabling dismissal of predictions with too low certainty. The model was trained and evaluated on a large, heterogeneous dataset containing AST results from blood infections collected during routine care across 30 European countries. Our results showed that the model could accurately predict susceptibility to a wide range of antibiotics, even when only a few AST results were included in the input. We also show that the model handles increasing information well, providing more accurate predictions as additional AST results become available. Finally, we demonstrate that predictions can be made within predefined confidence levels for most bacterial species and antibiotics, which allows control of both the major and very major error rates. We conclude that the combination of transformers and conditional inductive conformal predictions constitutes an appealing class of models for integrating and predicting diagnostic information.

## RESULTS

### A transformer-based model for antibiotic susceptibility prediction

We developed a transformer-based model to predict unavailable diagnostic information using multiple classifications. The input to the model is a set of words containing the available diagnostic information (results of antibiotic susceptibility testing; AST) and patient data (Figure 1). The AST results are assumed to be incomplete, where only a subset of the possible tests has available results. The model uses two transformers, which summarize the patient and AST information in two sentence-specific classification (CLS) vectors, one representing patient-specific information and one representing bacterial isolate-specific information. These vectors then serve as input to multiple antibiotic-specific feed-forward neural networks, each predicting the probability of susceptibility to the corresponding antibiotic. The uncertainty is estimated using inductive conformal prediction, conditioned on the susceptibility; hence, controlling the false-positive (major error) and false-negative (very major error) rates for each antibiotic (*34, 35*). The full details of the model architecture and the uncertainty estimation are provided in Methods.

**Figure 1.**
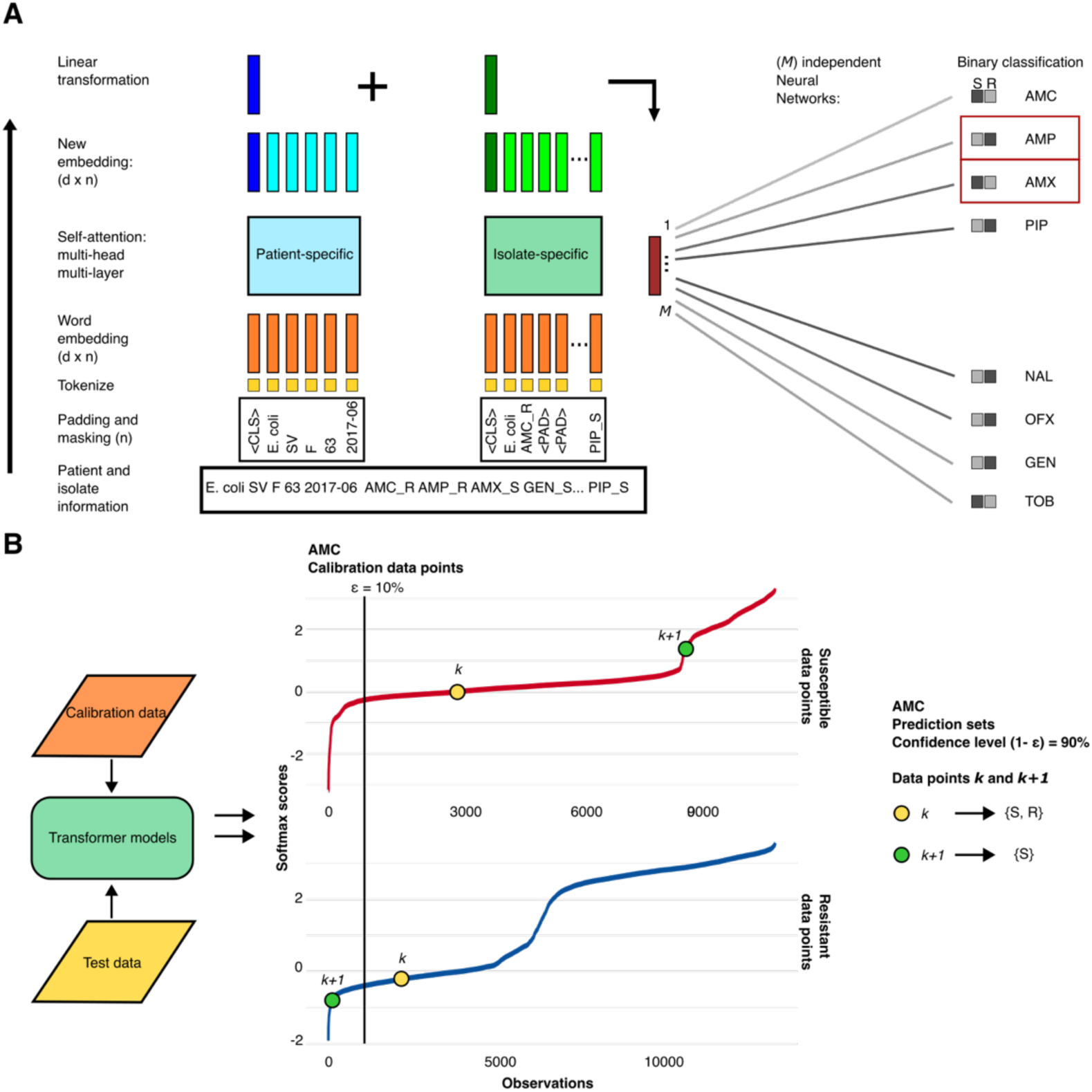
The proposed model architecture and uncertainty control. **(A)** The architecture of the proposed model. Both input sequences start with a classification word, *CLS*, followed by the bacterial species name. One of the input sequences also contains patient information (country, gender, age, and sampling date), while the other includes available antibiotic susceptibility testing (AST) data. Both input sequences are fixed to lengths L=5 for the patient information and L=21 for the AST data, padded with the *PAD* word. We use a linear embedding to represent the input words numerically, which are fed into the transformer encoders. The first vectors (*CLS*) from the outputs of each encoder are combined into a linear model and fed to *m* independent neural networks, each representing one antibiotic. The neural network outputs are two- dimensional vectors indicating susceptibility and resistance, respectively. **(B)** Uncertainty control. The neural network outputs undergo a softmax rescaling. A calibration dataset is used to build empirical distributions of conformity for resistant and susceptible predictions for each pathogen and antibiotic. The prediction regions for the data points 𝑘 and 𝑘 + 1 are built based on the deviation of the observed softmax score from the empirical distribution and the confidence interval threshold. See Methods for full details.

The model was trained and evaluated on data from The European Surveillance System (TESSy) (https://www.ecdc.europa.eu/en/publications-data/european-surveillance-system-tessy), which contains AST results for 1,161,303 *E. coli*, 301,506 *K. pneumoniae*, and 166,448 *P. aeruginosa* isolates collected from blood infections of patients in 30 European countries. The training and evaluation were performed on 20 (16, 18, and 11 for *E. coli*, *K. pneumoniae*, and *P. aeruginosa,* respectively) commonly used antibiotics that belong to five large and clinically relevant antibiotic classes: aminoglycosides, carbapenems, cephalosporins, penicillins, and quinolones (Table 1). A bacterial isolate was, on average, tested for susceptibility to 7.3, 8.8, and 7.8 antibiotics, with standard deviations of 1.9, 2.4, and 1.7 for *E. coli*, *K. pneumoniae*, and *P. aeruginosa*, respectively (Figures S1 and S2). The most commonly tested antibiotics were ceftazidime (CAZ) and ciprofloxacin (CIP), for which at least 93% of isolates were tested, regardless of pathogen, country of origin, or patient gender. In contrast, less than 10% of the *E. coli* and *K. pneumoniae* isolates were tested for piperacillin (PIP), moxifloxacin (MFX), and nalidixic acid (NAL, Figure S3). The susceptibility rate for *E. coli* isolates was lowest for the penicillins amoxicillin (AMX), ampicillin (AMP), and PIP (43%–50%), whereas for *K. pneumoniae* isolates it was lowest for PIP (20%). For other antibiotics and *P. aeruginosa* isolates, the susceptibility rates were higher, representing a more unbalanced dataset. A slightly higher susceptibility rate was observed among isolates collected from female patients (Figure S4).

**Table 1:** Bacterial isolates summary. The included antibiotics, the number of isolates from each species tested for each antibiotic, their resistance rates, and the weights used in the loss function.

| Antibiotic | *Escherichia coli* |  | *Klebsiella pneumoniae* |  | *Pseudomonas aeruginosa* |  | Loss function weights<br>(S/R)<sup>2</sup> |
| --- | --- | --- | --- | --- | --- | --- | --- |
|  | Isolates | %R<sup>1</sup> | Isolates | %R<sup>1</sup> | Isolates | %R<sup>1</sup> |  |
| AMC – Amoxicillin / clavulanic acid | 550 937 | 33% | 129 537 | 42% | – | – | 0.45/0.55 |
| AMP – Ampicillin | 800 283 | 57% | – | – | – | – | 0.45/0.55 |
| AMX – Amoxicillin | 293 369 | 55% | – | – | – | – | 0.45/0.55 |
| PIP – Piperacillin | 93 919 | 50% | 21 028 | 80% | 62 218 | 22% | 0.45/0.55 |
| TZP – Piperacillin/tazobactam | 646 892 | 8% | 164 551 | 29% | 152 233 | 16% | 0.15/85 |
| CAZ – Ceftazidime | 1 080 891 | 10% | 280 916 | 30% | 159 118 | 15% | 0.3/0.7 |
| CRO – Ceftriaxone | 350 947 | 11% | 90 851 | 30% | – | – | 0.3/0.7 |
| CTX – Cefotaxime | 990 234 | 11% | 253 971 | 30% | – | – | 0.3/0.7 |
| FEP – Cefepime | 336 526 | 10% | 102 213 | 33% | 65 619 | 16% | 0.3/0.7 |
| CIP – Ciprofloxacin | 1 117 333 | 22% | 286 899 | 31% | 160 669 | 20% | 0.3/0.7 |
| LVX – Levofloxacin | 291 403 | 23% | 73 574 | 29% | 45 636 | 24% | 0.3/0.7 |
| MXF – Moxifloxacin | 105 648 | 26% | 23 231 | 26% | – | – | 0.3/0.7 |
| NAL – Nalidixic acid | 63 468 | 27% | 16 532 | 36% | – | – | 0.45/0.55 |
| OFX – Ofloxacin | 136 704 | 19% | 33 764 | 36% | – | – | 0.3/0.7 |
| AMK – Amikacin | – | – | 202 968 | 9% | 130 264 | 9% | 0.15/85 |
| GEN – Gentamicin | 1 073 614 | 9% | 278 400 | 20% | 143 645 | 14% | 0.15/85 |
| TOB – Tobramycin | 564 295 | 10% | 145 914 | 27% | 119 954 | 13% | 0.15/85 |
| ETP – Ertapenem | – | – | 109 997 | 14% | – | – | 0.15/85 |
| IPM – Imipenem | – | – | 189 733 | 10% | 121 852 | 20% | 0.15/85 |
| MEM – Meropenem | – | – | 242 732 | 9% | 130 352 | 15% | 0.15/85 |

<sup>1</sup>Proportion of resistance bacterial isolates to each of the antibiotics.
<sup>2</sup>Weight on the cross-entropy loss function for the two different labels: susceptible/resistant.
Dash (–) indicates combinations of bacterial species and antibiotics that the model does not cover.

### Predictions of antibiotic susceptibility have high performance

To assess how well the model predicts the susceptibility of antibiotics not included as inputs, we used 70% of the bacterial isolates for 5-fold cross-validation and model training. We used 15% of the isolates for testing, and the remaining 15% for calibration of conformal prediction. During training, calibration, and testing, we selected a random number of antibiotics as input to the model (distribution available in Figure S5; mean 5.7, 6.3, and 5.7 with standard deviations of 1.3, 1.5, and 1.1 for *E. coli*, *K. pneumoniae*, and *P. aeruginosa* isolates, respectively; see Methods for details) together with patient data. The susceptibility of the remaining tested antibiotics was assumed to be unknown and, therefore, masked from the input data. The model’s predictions were then compared with the antibiotics masked from the input to evaluate performance. In this setting, the model had an overall high performance that did not differ substantially between folds in the cross-validation, with a standard error (se) below 0.015 for all antibiotics and bacterial species except the major error (ME) for the antibiotic PIP in *K. pneumoniae* (se = 0.016 for both training and test datasets). The performance was largely consistent across the training and test processes (Figures 2 and S6). There were, however, apparent differences in performance between the three bacterial species and between antibiotics. The F1 score (the harmonic mean of precision and recall) in the test dataset for *E. coli and K. pneumoniae* isolates was highest for cephalosporins (91% and 96% on average, respectively), and quinolones (86% and 93% on average, respectively), though lower for penicillins (74% and 88% on average, respectively, Figure 2A). For *P. aeruginosa* isolates, the F1 score was 84% for penicillins, and 82% for quinolones and cephalosporines. Carbapenems had an average F1 score of 90% and 76% for *K. pneumoniae* and *P. aeruginosa* isolates, respectively, while aminoglycosides had the lowest F1 scores: 65% for *E. coli* and 76% for *K. pneumoniae* and *P. aeruginosa* isolates.

**Fig. 2.**
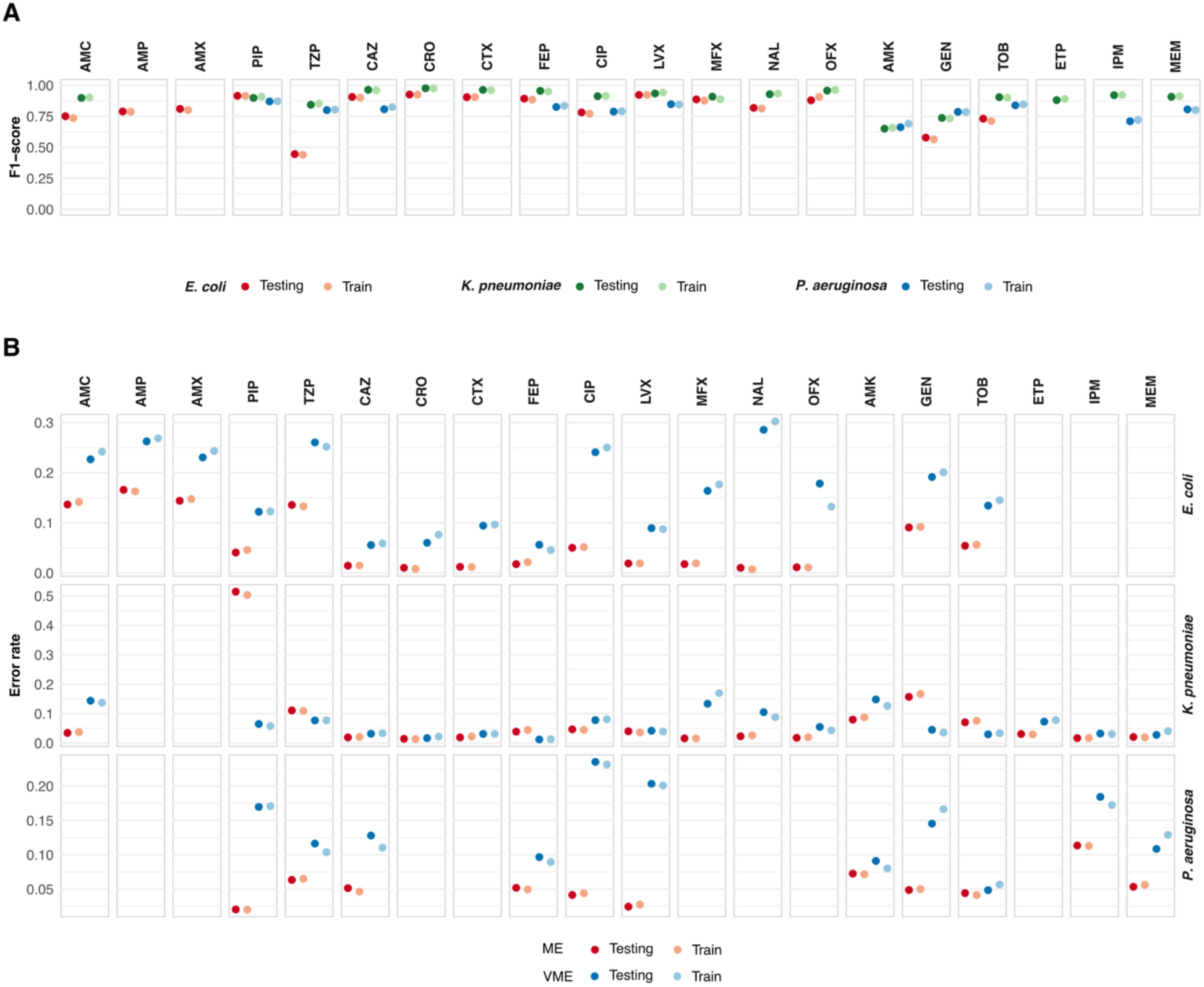
Performance of the transformer model. Results from the train and testing datasets: **(A)** F1-score, **(B)** major error (ME) rate, and very major error (VME) rate for the transformer model shown for each antibiotic and pathogen. Note that, due to data restrictions, not all antibiotics are assessed for all three pathogens.

Next, we evaluated the model based on the ME rate, defined as the proportion of susceptible bacterial isolates erroneously predicted as resistant, and the very major error (VME) rate, defined as the proportion of resistant isolates erroneously predicted as susceptible. The ME and VME rates are standard performance measures in antibiotic susceptibility testing, and are frequently used to evaluate and compare diagnostic methods (*36*). Based on the test dataset, cephalosporins had, on average over all predictions in the test data, an ME rate of 1.4% (1%– 1.8%), 2.3% (1.5%–3.9%), and 5.2% (5.1%–5.2%) for *E. coli*, *K. pneumoniae*, and *P. aeruginosa* isolates, respectively, while the corresponding VME rate was, on average, 6.7% (5.6%–9.4%), 2.3% (1.2%–3.2%), and 12.2% (9.7%–12.8%) (Figure 2B). For quinolones, the average ME rate was 2.2% (2%–5%), 2.9% (1.6%–4.7%), and 3.3% (2.5%–4.2%), while the VME rate was, on average 19.2% (8.9%–28.6%), 8.3% (4.2%–13.4%), and 21.9% (20.3%–23.5%) for *E. coli*, *K. pneumoniae*, and *P. aeruginosa* isolates, respectively. For *E. coli* isolates, the penicillins had a higher overall average ME rate of 12.5% (4.1%–16.6%). Here, PIP had the lowest VME rate (12.2%), while the other penicillins had rates between 23% and 26%. For *K. pneumoniae* and *P. aeruginosa* isolates, the average ME for penicillins, excluding PIP, was 7.3% and 6.3%, respectively, while the VME was 11%. We noted that the model struggled to predict PIP resistance in *K. pneumoniae* with an ME as high as 51%. This may be a consequence of data imbalance, where *K. pneumoniae* had a resistance rate of 80% to PIP (compared to 50% and 22% for *E. coli* and *P. aeruginosa*, respectively) but accounted for only 11% of the data for this antibiotic. Aminoglycosides had average ME/VME rates of 7.3/16.3, 10.2/7.4, and 5.5/9.5, for *E. coli, K. pneumoniae*, and *P. aeruginosa* isolates, respectively. Finally, the ME and VME rates for carbapenems for *K. pneumoniae* isolates were 2.3% (1.7%–3.1%) and 4.4% (2.8%–7.3%), respectively, and for *P. aeruginosa* isolates, 8.4% (5.3%–11.4%) and 14.6% (10.9%–18.4%), respectively.

The number of available AST results influenced the model’s performance. When the number of AST results used as input to the model increased from four to eight antibiotics, significant reductions were seen in the ME rate for penicillins, where AMX and AMP dropped from 34% and 22% to 3% and 7%, respectively, and the VME of CIP, which fell from 36% to 7% in *E. coli* isolates (Figure 3D). Interestingly, the drop for other antibiotics and pathogens was more modest or non-existent. A substantial reduction in the VME rate was also observed across most antibiotics and pathogens.

**Fig. 3.**
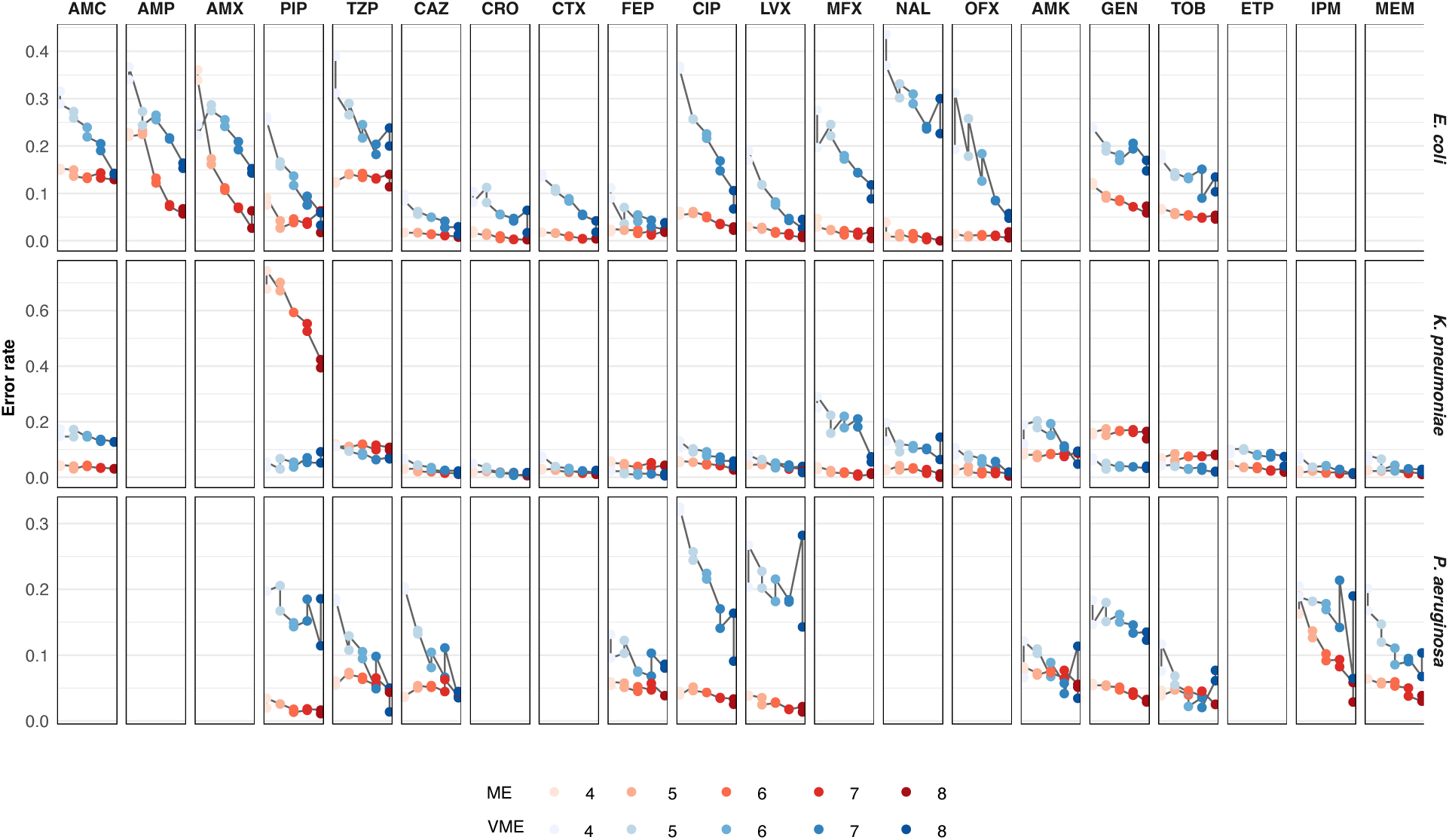
The predictive performance of the model as a function of the number of antibiotic susceptibility testing (AST) results included in the input. Major error (ME) and very major error (VME) rates from light to dark: 4 to 8 AST results are shown for each antibiotic and pathogen. Note that, due to data restrictions, not all antibiotics are assessed for all three pathogens.

When eight AST results were included in the input, fourteen of sixteen antibiotics in *E. coli* isolates, sixteen of twenty antibiotics in *K. pneumoniae*, and all antibiotics in *P. aeruginosa* had an ME rate below 10%, and below 5% for eleven, thirteen, and ten antibiotics in each pathogens, respectively. Similarly, the VME rate was below 10% for nine, sixteen, and ten, and below 5% for six, twelve, and three of the antibiotics corresponding to each pathogen. Going from four to eight input AST results increased the F1 scores from 75% to 87% for *E. coli*, from 85% to 92% for *K. pneumoniae*, and from 75% to 84% for *P. aeruginosa*, on average. The model’s overall high performance, as indicated by its F1 score, ME, and VME rates, suggests that it can be used to predict complete susceptibility patterns for bacterial isolates accurately.

### Control of the major and very major error rates at the patient level

In clinical practice, decisions are taken at the patient level. It is thus vital that decision support based on predictions also conveys information about the confidence in the prediction.

Indeed, if the uncertainty is too high, the prediction may need to be considered with care or completely dismissed until more diagnostic information becomes available. Therefore, we implemented an algorithm that assigns a quantitative measure of uncertainty to each individual prediction (*34*). The algorithm was based on conditional inductive conformal prediction, where the certainty of each prediction was derived from a conformity measure, defined in our case as the softmax score from each antibiotic-specific neural network (Figure 1B). The distributions of softmax scores for both susceptible and resistant bacterial isolates were empirically calculated for each antibiotic and each of the three pathogens, using a dedicated calibration dataset. These distributions were used to determine whether a new prediction aligns with the susceptible and/or resistant isolates in the calibration dataset and, based on a predefined confidence level, is deemed sufficiently certain. The output of a final prediction for one antibiotic is a prediction set, with either a) a single label, i.e., either susceptible or resistant, when there is enough conformity to only one of the softmax distributions; b) multiple labels, i.e. both susceptible and resistant, if there is enough conformity for both softmax distributions; or c) no label if there is no conformity to either group. The proportion of predictions that will, on average, have the correct label in their prediction set is governed by the confidence level 1 − 𝜀. Note that in this setting, 𝜀 corresponds to the average ME and the VME rates for susceptible and resistant bacterial isolates for each pathogen, respectively (see Methods for full details).

The empirically derived ME and VME rates were close to the prespecified values of ε for all antibiotics (Figures S7-S11). For example, at a confidence level of 90% (𝜀 = 0.1), the observed ME rates were, on average, 10.1%, 9.9%, and 9.9% (standard deviation 0.4%, 1.3%, and 0.3%), and the observed VME rates were, on average 9.7%, 9.4%, and 9.3% (standard deviation 1.43%, 0.95%, and 1.7%) for *E. coli*, *K. pneumoniae*, and *P. aeruginosa* isolates, respectively. The concordance between pre-specified and observed error rates was sustained at higher confidence levels where, for a confidence level of 95%, the average ME rates were 5.1%, 5.1%, and 5% (standard deviation 0.5%, 1%, and 0.3%) and the average VME rates were 5%, 4.7%, and 5% (standard deviation 0.8%, 0.6%, and 1.4%) for *E. coli*, *K. pneumoniae*, and *P. aeruginosa* isolates, respectively. Thus, the results showed that the specified confidence levels calculated from empirical distributions were sufficiently stable between datasets.

### The model could confidently predict the phenotype for a large majority of bacterial isolates and antibiotics

At a confidence level of 90%, as many as 82%, 89%, and 86% of the predictions were unambiguous and correct (only the correct label was in the prediction set) for *E. coli*, *K. pneumoniae*, and *P. aeruginosa* isolates, respectively; however, this number varied between antibiotics (Figure 4A, 5A, 6A). Cephalosporins had a high proportion of correct unambiguous predictions (90%) across the three pathogens, and quinolones had 90% for *K. pneumoniae*. For the remaining antibiotics, between 82% and 89% of the predictions were correct and unambiguous, except for penicillins for *E. coli* (71%) and quinolones for *P. aeruginosa*, which showed higher uncertainty. The proportion of unambiguous predictions was, as expected, reduced when the confidence level was increased; to 78%, 89%, and 77% at a 95% confidence level and, finally, to 72%, 85%, and 63% at a 97.5% confidence level for *E. coli*, *K. pneumoniae*, and *P. aeruginosa*, respectively. On the other hand, the number of unambiguous predictions increased as more diagnostic information was provided to the model. At a 90% confidence level, the proportion of correct unambiguous predictions increased from, on average, 76%, 84%, and 83% when four AST results were included in the input, to 89%, 91%, and 91% when eight AST results were included in the input for *E. coli, K. pneumoniae*, and *P. aeruginosa* isolates, respectively (Figure 4B, 5B, 6B). The increase was substantial for predicting susceptibility to the penicillins AMP and AMX in *E. coli* isolates, to PIP and OFX in *K. pneumoniae* isolates, and for predicting susceptibility and resistance to CIP, as well as susceptibility to LVX and IPM in *P. aeruginosa* isolates. With eight input AST results, the model achieved 90% correct, unambiguous predictions for most of the remaining masked antibiotics. This was also true for higher confidence levels, where the proportion of unambiguous and correct predictions with eight input AST results was 89% and 84% for 95% and 97.5% confidence, respectively.

**Fig. 4.**
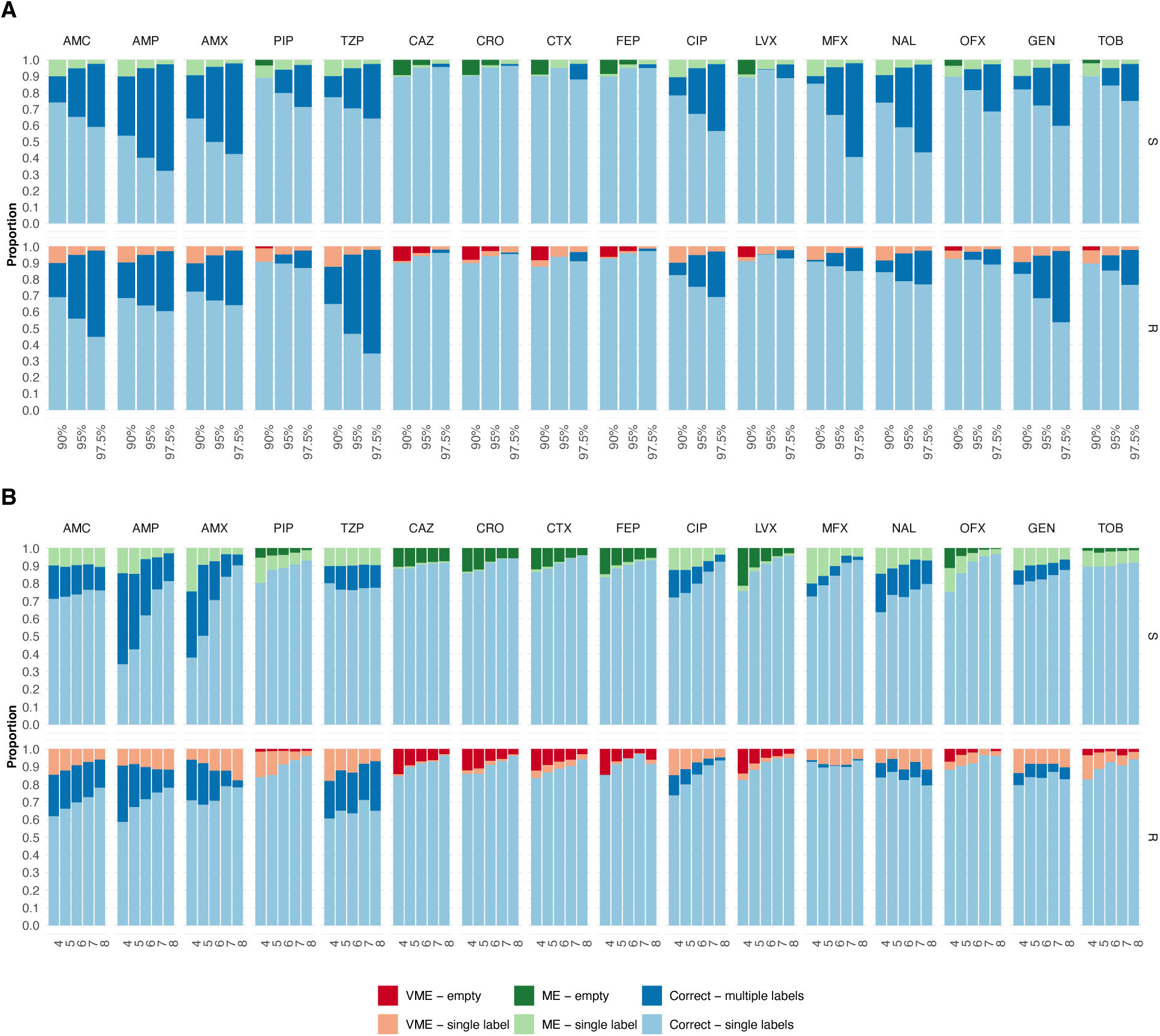
The proportion of correct predictions with respect to the number of labels in the prediction set for *E. coli* isolates. The proportion of correct predictions with a single label (light) and multiple labels (dark), major errors (MEs), and very major errors (VMEs) with a single label (light) and empty set (dark) predictions for resistant (R) and susceptible (S) isolates, shown for each antibiotic and pathogen using three different confidence levels: 90%, 95%, and 97.5%. **(A)** The proportions are shown for each antibiotic at three confidence levels (90%, 95%, and 97.5%), considering all possible numbers of input AST results. **(B)** The proportions are shown as a function of the number of input AST results (90% confidence level).

**Fig. 5.**
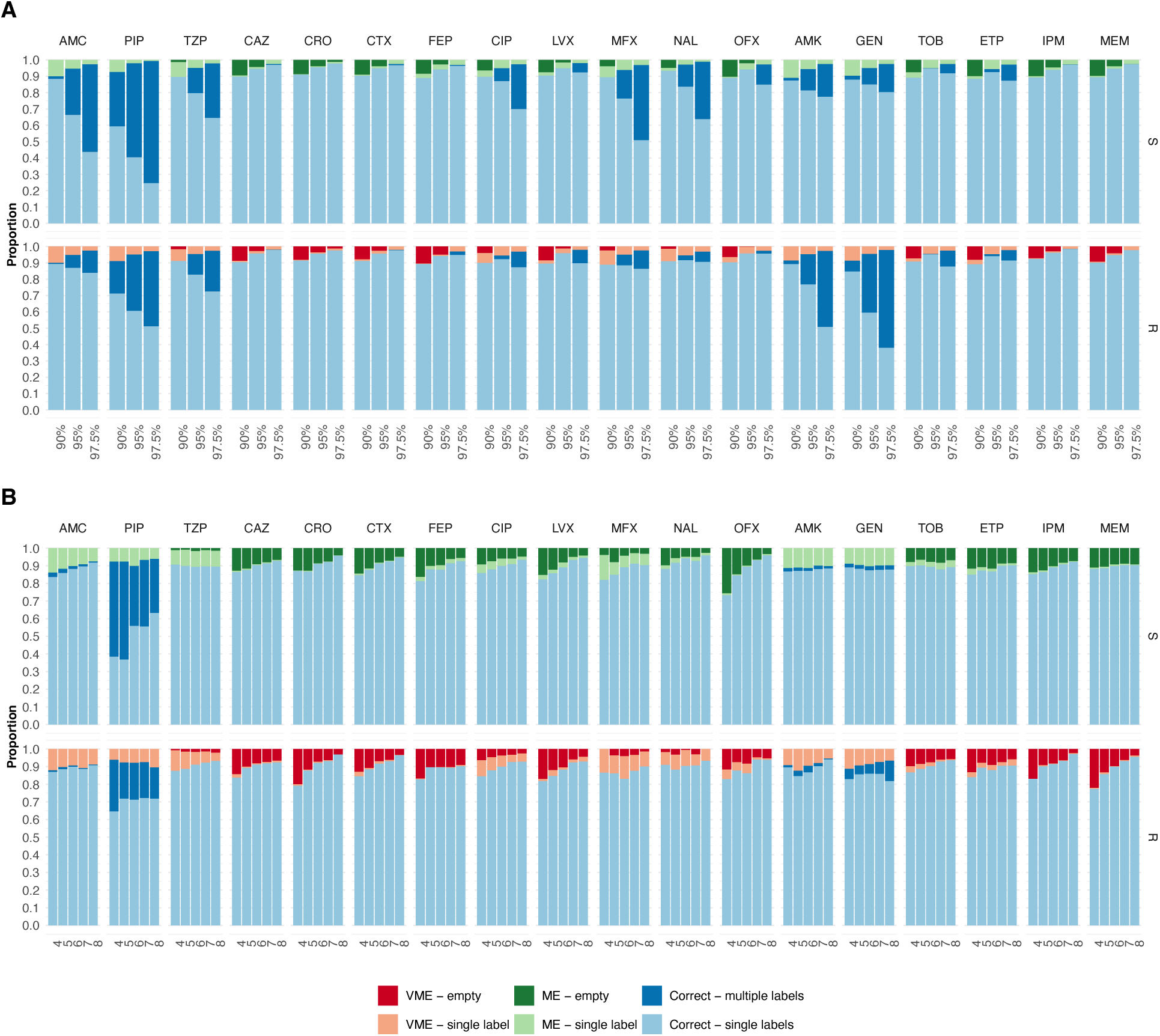
The proportion of correct predictions with respect to the number of labels in the prediction set for *K. pneumoniae* isolates. The proportion of correct predictions with a single label (light) and multiple labels (dark), major errors (MEs), and very major errors (VMEs) with a single label (light) and empty set (dark) predictions for resistant (R) and susceptible (S) isolates, shown for each antibiotic and pathogen using three different confidence levels: 90%, 95%, and 97.5%. **(A)** The proportions are shown for each antibiotic at three confidence levels (90%, 95%, and 97.5%), considering all possible numbers of input AST results. **(B)** The proportions are shown as a function of the number of input AST results (90% confidence level).

**Fig. 6.**
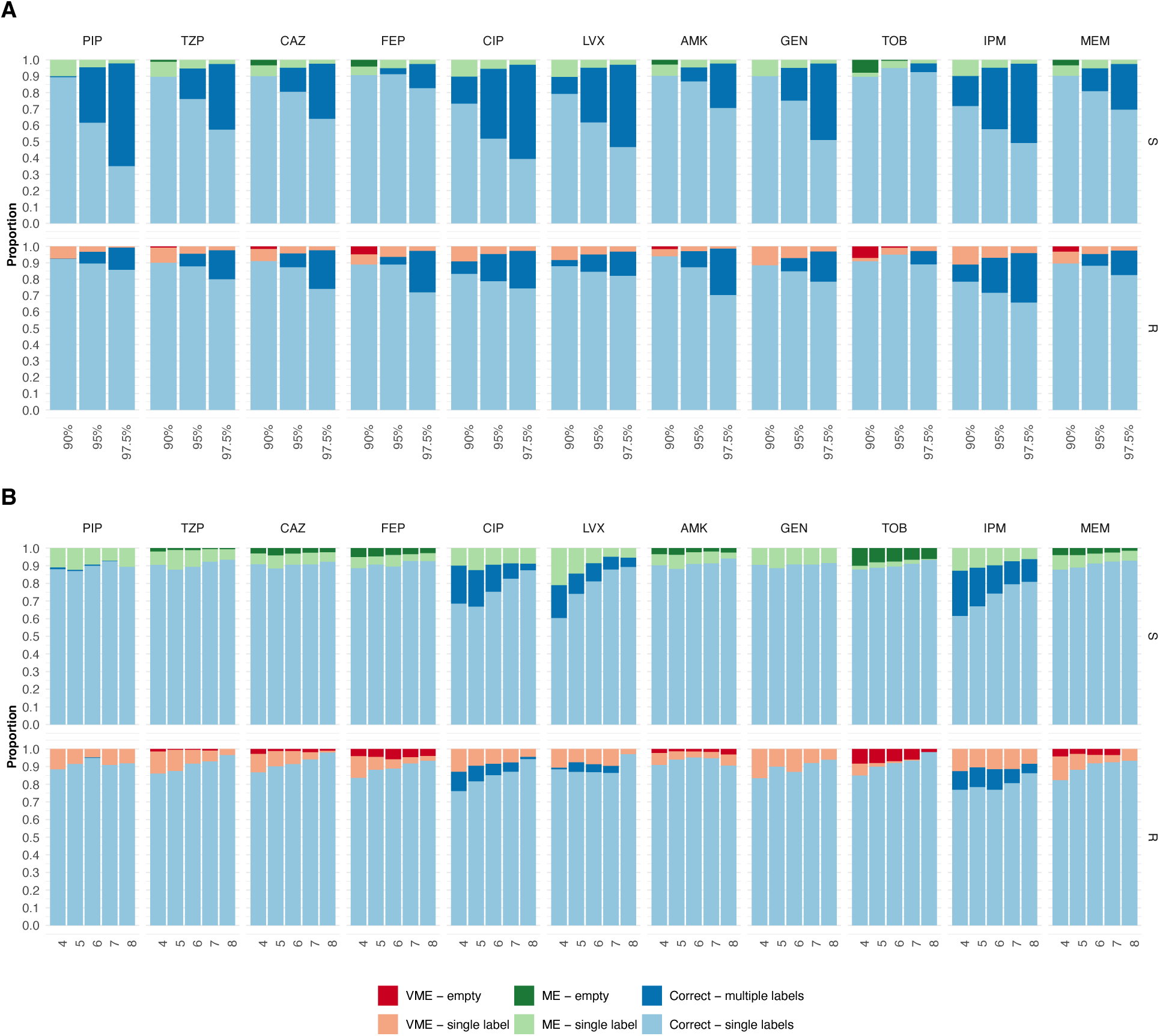
The proportion of correct predictions with respect to the number of labels in the prediction set for *P. aeruginosa* isolates. The proportion of correct predictions with a single label (light) and multiple labels (dark), major errors (MEs), and very major errors (VMEs) with a single label (light) and empty set (dark) predictions for resistant (R) and susceptible (S) isolates, shown for each antibiotic and pathogen using three different confidence levels: 90%, 95%, and 97.5%. **(A)** The proportions are shown for each antibiotic at three confidence levels (90%, 95%, and 97.5%), considering all possible numbers of input AST results. **(B)** The proportions are shown as a function of the number of input AST results (90% confidence level).

## DISCUSSION

In this study, we present a method that uses a transformer model to predict unavailable diagnostic information. When combined with conditional inductive conformal prediction to estimate the predictive uncertainty at the patient level, the method can abstain from making decisions unless the confidence is deemed sufficiently high. The model was applied to the diagnostics of infectious bacteria, a field facing rapidly rising societal and economic costs due to the growing challenges related to antibiotic resistance (*37, 38*). We showed that only a few antibiotic susceptibility test (AST) results are sufficient to accurately derive a more complete resistance profile. This has the potential to guide antibiotic therapy for serious infections, where treatment must begin before all diagnostic tests are completed.

The method was trained on a large, heterogeneous dataset comprising over 12 million AST results from more than a million bacterial isolates of three pathogens collected from 30 European countries. This dataset was compiled for surveillance purposes and contains AST results originally produced in routine diagnostics. Validation, in which AST results were randomly excluded, showed that the model had generally high accuracy in predicting antibiotic susceptibility. The performance improved further as more ASTs were included in the input data, demonstrating that the model could efficiently incorporate diagnostic information as it becomes available and, thus, over time, produce more certain predictions. Indeed, when AST for eight antibiotics was used as input, the model could predict most susceptibilities with a VME rate (false-negative rate) as low as 10%. This indicates that AI prediction can serve as a viable complement to laboratory diagnostic tests and may be utilized to save time, alleviate suffering, and reduce economic costs.

Providing information about predictive uncertainty is crucial in diagnostics, where population-level knowledge is used to predict outcomes for individual patients. We addressed this challenge by implementing an algorithm based on conditional inductive conformal prediction (*34, 39, 40*) to provide each prediction with an accompanying confidence score. In our application, the predefined confidence corresponded to ME and VME rates, yielding predictions that limit false-positive and false-negative rates to the desired levels. During model testing, we found consistent error rates across datasets and, for cephalosporins and quinolones, the prediction sets mainly contained a single, correct label. However, a higher variation was observed for penicillins, resulting in a larger proportion of ambiguous predictions (multiple labels). Furthermore, our implementation separates the uncertainty for susceptible and resistant predictions. This is valuable in the diagnostics of bacterial infections, where a VME, i.e., the incorrect identification of a resistant isolate, may lead to ineffective antibiotic treatment.

Therefore, VMEs are often considered to be the most serious errors, especially for life-threatening infections. The ability to set confidence scores for susceptible and resistant predictions individually enables the model to be adjusted for different clinical scenarios.

The model showed a clear difference in predictive performance between antibiotics, notoriously lower for penicillins, especially compared to cephalosporins. Historically, penicillins were among the earliest classes of antibiotics and the first beta-lactam antibiotics introduced.

Resistance to beta-lactam antibiotics is typically caused by enzymes that hydrolyze the antibiotics (*41*). Penicillins have been widely used for over 80 years, and bacteria can consequently acquire a wide diversity of resistance genes (*42*). In contrast, resistance to cephalosporins – a later generation of beta-lactam antibiotics – is, to a larger extent, dependent on additional genetic events, such as mutations in chromosomal genes or the acquisition of broader-spectrum resistance mechanisms (*41*). These patterns were reflected in the data, where resistance to penicillins was common. Indeed, as many as 87.7% of the *E. coli* and 61% of the *K. pneumoniae* bacterial isolates resistant to a single antibiotic were resistant to a penicillin.

Furthermore, it is plausible that the order of evolution of multidrug-resistant bacteria affects the performance of our model. We also consider resistance phenotypes that are initially acquired to be harder to predict than those that commonly appear in later stages of evolution, simply because there are no other correlating susceptibilities. Results from additional diagnostic assays, such as targeted molecular tests that detect antibiotic resistance genes and genomic data, could further improve performance. Indeed, the flexibility of the transformers allows incorporating other types of diagnostic information, including genotypic information, as new words in the input sentence.

Ultimately, the AI methodology presented here has the potential to enhance the diagnosis of infections caused by antibiotic-resistant bacteria. We argue that data-driven methods have the potential to replace selected diagnostic assays, thereby providing physicians with more comprehensive decision support at an earlier stage. This has the potential to improve the treatment of bacterial infections, thereby decreasing patient morbidity and mortality, reducing costs, and limiting the overprescription of antibiotics.

## MATERIALS AND METHODS

### Data description

This study is based on data from the European Surveillance System (TESSy), collected as part of surveillance conducted by the European Centre for Disease Prevention and Control. The dataset contains more than 12 million antibiotic susceptibility testing (AST) results for bacteria isolated from blood and cerebrospinal fluid of hospitalized patients across 30 European countries. For each bacterial isolate, we retrieved the species name, AST results, the patient’s gender and age, the date of bacterial isolation, and the country where the test was performed.

This metadata will be referred to collectively as the patient data. The analysis was limited to the susceptibility of *Escherichia coli, Klebsiella pneumoniae,* and *Pseudomonas aeruginosa* isolates, which were the most common species, from blood infections collected between 2007 and 2020. Tests that resulted in either a resistant (R) or a susceptible (S) result were included, while tests with an I result were excluded. Only antibiotics with a resistance rate of at least 8% were considered. This resulted in data covering sixteen antibiotics from five clinically relevant classes: aminoglycosides, cephalosporins, penicillins, quinolones, and carbapenems (Table 1).

Furthermore, isolates tested for fewer than five antibiotics were removed. If an antibiotic was tested multiple times for the same bacterial isolate, only the most recent test was included. The final dataset contained 𝑛 = 1,629,257 bacterial isolates with 12,433,856 AST results, resulting in an average of 7.6 tests (standard deviation of 2.1) per isolate. After randomization, the bacterial isolates were divided into three datasets: a training dataset (70%), a calibration dataset (15%), and a test dataset (15%). The train dataset was further divided into five groups for 5-fold cross-validation.

### Data expansion

To increase the number of AST result combinations, each dataset was expanded. For each bacterial isolate 𝑗, multiple data points were generated by randomly splitting the susceptibility test results into two groups. The first group, together with the patent information, 𝑥_!_, was considered known and, thus, used as input to the model. The second group, 𝑦_!_, was hidden from the model and used to evaluate the predictive performance. The full details of the data expansion are provided in Supplemental Materials (Table S1). A made-up example of the information available for a bacterial isolate is presented below:

*Available information for bacterial isolate* 𝑗: “ESCCOL SV 30 M 2013_01 LVX_R AMC_S AMP_S TZP_R CTX_S GEN_S”, represents a. *E. coli* bacterium isolated at a Swedish hospital (SV) from a 30-year-old (30) male (M) patient in January 2013 (2013_01) where the isolated bacterium was tested against six antibiotics. The AST results indicated resistance to the antibiotics levofloxacin (LVX) and piperacillin/tazobactam (TZP), and susceptibility to amoxicillin/clavulanic acid (AMC), ampicillin (AMP), cefotaxime (CTX), and gentamicin (GEN). Two example data points, (𝑥_"_, 𝑦_"_) and (𝑥_"#_, 𝑦_"#_), that could, potentially, be created from this isolate: (𝑥_"_, 𝑦_"_) = (“ESCCOL SV 30 M 2013_01 LVX_R AMC_S AMP_S TZP_R”, “CTX_S GEN_S”), (𝑥_"#_, 𝑦_"#_) = (“ESCCOL SV 30 M 2013_01 LVX_R AMC_S AMP_S CTX_S GEN_S”, “TZP_R”).

### Model Architecture and Training

Given a data point, (𝑥_"_, 𝑦_"_), the model takes the input 𝑥_"_ and makes predictions 𝑦G_"_ of the susceptibilities 𝑦_"_. The input sentence 𝑥_"_ is split into patient data and AST data, both of which contain the word for the bacterial species, and both are complemented at the start with the classification word, *CLS.* The patient data and the AST data are padded to a length 𝐿 = 6 and 𝐿 = 21 with *PAD* words if needed. Each word is then converted into a linear embedding representation in the form of a 𝑑-dimensional vector that provides semantic meaning to the model (𝑑 = 128). These word embeddings are passed through two transformer encoder layers, each with two attention heads, followed by an add-and-normalize layer, a position-wise feed- forward layer (using 256 nodes), and, finally, another add-and-normalize layer. The first vector of the output from each encoder– representing the *CLS* word and containing information at the sentence level– are concatenated and fed to a linear transformation. The output is used as input to 𝑀 = 20 independent antibiotic-specific feed-forward networks, each of depth two. The intermediate layer of the network has 128 nodes with rectified linear unit (ReLU) activation function, followed by a normalization step, while the final layer outputs a linear transformation to vectors of length two, which are used for binary classification. The isolate was classified as resistant or susceptible based on the largest output value. Additionally, models with the combinations of 1 or 2 encoder heads and layers, with word embedding of size 64 or 128, and a position-wise feed-forward layer of 128 or 256, with patient data and no patient data, were trained (Figure S13-S15). The proposed model achieved higher accuracy, lower ME, and lower VME in 23, 14, and 10 out of 45 pathogen-antibiotic combinations, respectively. The combination of one head, one layer, with word embedding of size 128 and a position-wise feed- forward layer of size 256 was the only one to have more pathogen-antibiotic combinations with lower VME (13).

The model was trained as follows. At each epoch, 248,000 bacterial isolates were randomly sampled from the train dataset and expanded as described in Methods – Data expansion. The model was then trained on 512,000 randomly selected data points from the data expansion step, divided into mini-batches of size 512. The loss was based on the cross-entropy between the known (𝑦_"_) and predicted (𝑦G_"_) labels. The Adam optimizer was used to minimize the loss over 700 epochs with a fixed learning rate of 1x10^-6^. The model was implemented and trained using Pytorch version 1.7.1.

### Uncertainty control

An algorithm based on conditional inductive conformal prediction was used to quantify the uncertainty of the predictions (*34*) with respect to the antibiotic and label (i.e., “susceptible” and “resistant”) (34). Conformal prediction uses empirical data to estimate uncertainty and has previously been shown to suit complex diagnostic data for which valid distributional assumptions may be hard to make (*43*). For a data point (𝑥_"_, 𝑦_"_) belonging to pathogen 𝑞, the algorithm estimates a prediction set Γ^$,&^, containing the predictions that are deemed sufficiently confident given a predefined confidence level 1 − 𝜀. The uncertainty for a prediction was based on its conformity measure, defined as the softmax transformation of the outputs of a neural network. The conformity measure and, thus, the uncertainty, was derived individually for each antibiotic and each label (i.e., resistance or susceptible).

We estimated the empirical distributions for each conformity measure from the calibration dataset separately for each pathogen, which, for an antibiotic 𝑎 and pathogen 𝑞, were assumed to contain 𝑙_&,’_= 𝑙_&,’,(_ + 𝑙_&,’,)_data points, where 𝑙_&,’,(_ and 𝑙_&,’,)_are the number of data points for susceptible and resistant bacterial isolates of pathogen 𝑞, respectively. For a data point (𝑥_"_, 𝑦_"_) derived from pathogen 𝑞, let 𝛼^’,(^ and 𝛼^’,)^ denote the softmax score for the prediction of susceptibility and resistance to antibiotic 𝑎, respectively. The prediction sets were decided based on the empirical p-values 𝑝^&,’,(^ and 𝑝^&,’,)^, which were calculated according to

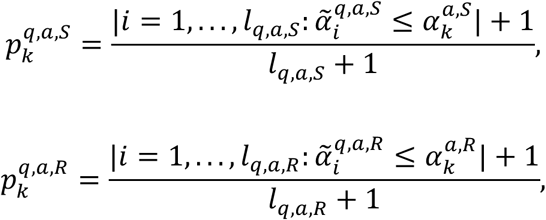

where 𝛼V ^&,’,(^ and 𝛼V^&,’,)^ denotes the softmax scores calculated using the data points in the calibration dataset for isolates from pathogen 𝑞. At a confidence 1 − 𝜀, the prediction set was then formed by

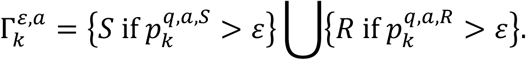

### Performance

The model’s predictive performance was computed based on 257,784, 242,121, and 114,410 train; 285,897, 223,661, and 97,542 calibration; and 280,428, 223,665, and 100,927 test individual AST results for *E. coli*, *K. pneumoniae*, and *P. aeruginosa* isolates, respectively. To evaluate the overall model performance, the F1 score was calculated. In addition, the *major error* (ME) rate, defined as the proportion of true susceptible bacterial isolates erroneously predicted as resistant, and the *very major error* (VME) rate, defined as the proportion of true resistant isolates erroneously predicted as susceptible, were also calculated. To measure the performance of the uncertainty control, true predictions were defined as prediction regions containing the true label for each antibiotic, and false predictions were those containing either no labels or only the wrong one.

For comparison, the following naive classifier was included (Figure S7). For each of the bacterial species, the frequencies between all pairwise antibiotics and their susceptibility profile were calculated in the training dataset. Then, the ratios 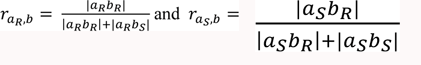, between antibiotic 𝑎, with resistance (𝑅) and susceptible (𝑆) profiles, and the antibiotic 𝑏, with resistance profile, were calculated for all pairwise combinations of antibiotics. Finally, for a new prediction of antibiotic 𝑏 in 𝑦_"_, resistance was deemed if the average 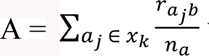 was larger than 0.5, where 𝑎 represent the antibiotics in 𝑥 with resistance 𝑗 ∈ {𝑅, 𝑆}, and 𝑛_’_is the number of antibiotics in 𝑥_"_, the isolate was deemed susceptible otherwise. To investigate the effect of patient data, we compared our results with a model based solely on AST results (Figure S6). The VMEs for the three bacterial species were found to be significantly smaller (p-val<0.01) when the original model was compared to the model without patient data and to the naive classifier, as determined by Wilcoxon signed-rank tests.

## ACKNOWLEDGMENTS

Data from The European Surveillance System – TESSy between 2007 to 2020, provided by Andorra, Albania, Armenia, Austria, Azerbaijan, Bosnia and Herzegovina, Belgium, Bulgaria, Belarus, Switzerland, Cyprus, Czechia, Germany, Denmark, Estonia, Greece, Spain, Finland, France, Georgia, Croatia, Hungary, Ireland, Israel, Iceland, Italy, Kyrgyzstan, Kazakhstan, Liechtenstein, Lithuania, Luxembourg, Latvia, Monaco, Republic of Moldova, Montenegro, Republic of North Macedonia, Malta, Netherlands, Norway, Poland, Portugal, Romania, Serbia, Russian Federation, Sweden, Slovenia, Slovakia, San Marino, Tajikistan, Turkmenistan, Turkey, Ukraine, United Kingdom, Uzbekistan, Kosovo and released by The European Centre for Disease Prevention and Control (ECDC). The views and opinions of the authors expressed herein do not necessarily state or reflect those of ECDC. The accuracy of the authors’ statistical analysis and the findings they report are not the responsibility of ECDC. ECDC is not responsible for conclusions or opinions drawn from the data provided. ECDC is not responsible for the correctness of the data and data management, data merging, and data collation after the provision of the data. ECDC shall not be held liable for improper or incorrect use of the data.

## DATA AVAILABILITY STATEMENT

The data used in this study belongs to the European Surveillance System – TESSy. We acknowledge the European Surveillance System – TESSy for data availability. The model is available in the GitHub repository https://github.com/indajuan/Confidence-based-Prediction-of-Antibiotic-Resistance.

## FUNDING

Swedish Research Council (VR), grant 2018-02835 (EK) Swedish Research Council (VR), grant 2019-03482 (EK) Chalmers AI Research Centre (CHAIR), (EK) National Health and Medical Research Council of Australia under the framework of JPI AMR, grant 2031902, SEARCHER

## CONFLICTS OF INTEREST

The authors declare no conflict of interest.

## SUPPLEMENTAL MATERIALS

Figure S1. The number of susceptible (opaque) and resistant (transparent) bacterial isolates tested for each antibiotic and pathogen in the dataset.

Figure S2. Distribution of the number of antibiotics tested per bacterial isolate for each pathogen in the dataset.

Figure S3. Distribution of the proportion of bacterial isolates tested against each antibiotic for female (opaque) and male (transparent) patients and for each pathogen across countries in the dataset. The center line, box limits, and whiskers represent the median, upper, and lower quartiles, and 1.5 times the interquartile range, respectively.

Figure S4. The proportion of bacterial isolates susceptible to each antibiotic for female (opaque) and male (transparent) patients and for each pathogen across countries in the dataset. The center line, box limits, and whiskers represent the median, upper, and lower quartiles, and 1.5 times the interquartile range, respectively.

Figure S5. Histogram of the number of antibiotic susceptibility testing (AST) results used as input for the model during training at each epoch for each pathogen.

Figure S6. Performance of the cross-validation. The major error (red) and very major error (VME) rates for each fold are represented by dots, the horizontal line, box limits, and whiskers represent the median, upper, and lower quartiles, and 1.5 times the interquartile range, respectively, across folds.

Figure S7. Performance of the model with and without patient data and the naive classifier.

Figure S8-12. Empirical and expected error rates and performance of the model after conditional inductive conformal prediction. For each pathogen and penicillins (S8), cephalosporins (S9), quinolones (S10), aminoglycosides (S11), and carbapenems (S12) antibiotics.

Figure S13-S15. Performance of the model with different hyperparameters: 1 or 2 heads (1H, 2H), 1 or 2 layers (1L, 2L), word embedding of size 64 and position-wise feed-forward layer of 128 (small), or word embedding of size 128 and position-wise feed-forward layer of 256 (large), with patient and AST data, or without patient data (no patient), for (S13) E. coli, (S14) K. pneumoniae, and (S15) P. aeruginosa.

Table S1. Data expansion. For isolate *j* with 𝐴_!_ AST results, we created a fixed number of data points at each epoch of training, calibration, and testing. The number of AST results, 𝑎_"_, included in the input sentence 𝑥_"_ of data point 𝑘 was sampled from a binomial distribution with parameters 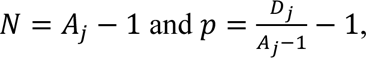, listed below. If 𝑎 < 4, the data point was dismissed. The number of antibiotic susceptibilities to predict for data point 𝑘 was given by 𝐴_!_ − 𝑎_"_, and the true susceptibility test results are summarized in the sequence 𝑦_"_. The decision on which antibiotics were included in 𝑥_"_ and 𝑦_"_ was done randomly for all data points.

Table S2. Performance of the transformer model. Results from the test dataset: F1-score, major error (ME) rate, and very major error (VME) rate shown for each antibiotic and pathogen, considering all possible numbers of input AST results. Dash (–) indicates combinations of bacterial species and antibiotics that the model does not cover.

Table S3. Performance of the transformer model as a function of the number of antibiotic susceptibility testing (AST) results included in the input. Results from the test dataset: F1-score, major error (ME) rate, and very major error (VME) rate shown for each antibiotic, pathogen, and the number of antibiotic susceptibility testing (AST) results included in the input. Dash (–) indicates combinations of bacterial species and antibiotics that the model does not cover. We limited the table to 8 AST results in the input due to the low number of isolates tested for 10 or more antibiotics in the data.

## Notes

https://github.com/indajuan/Confidence-based-Prediction-of-Antibiotic-Resistance

### Competing Interest Statement

The authors have declared no competing interest.

### Summary of Updates

The dataset was expanded. The model includes three different bacterial species.

