## Supplemental materials for "Confidence-based Prediction of Antibiotic Resistance at the Patient Level"

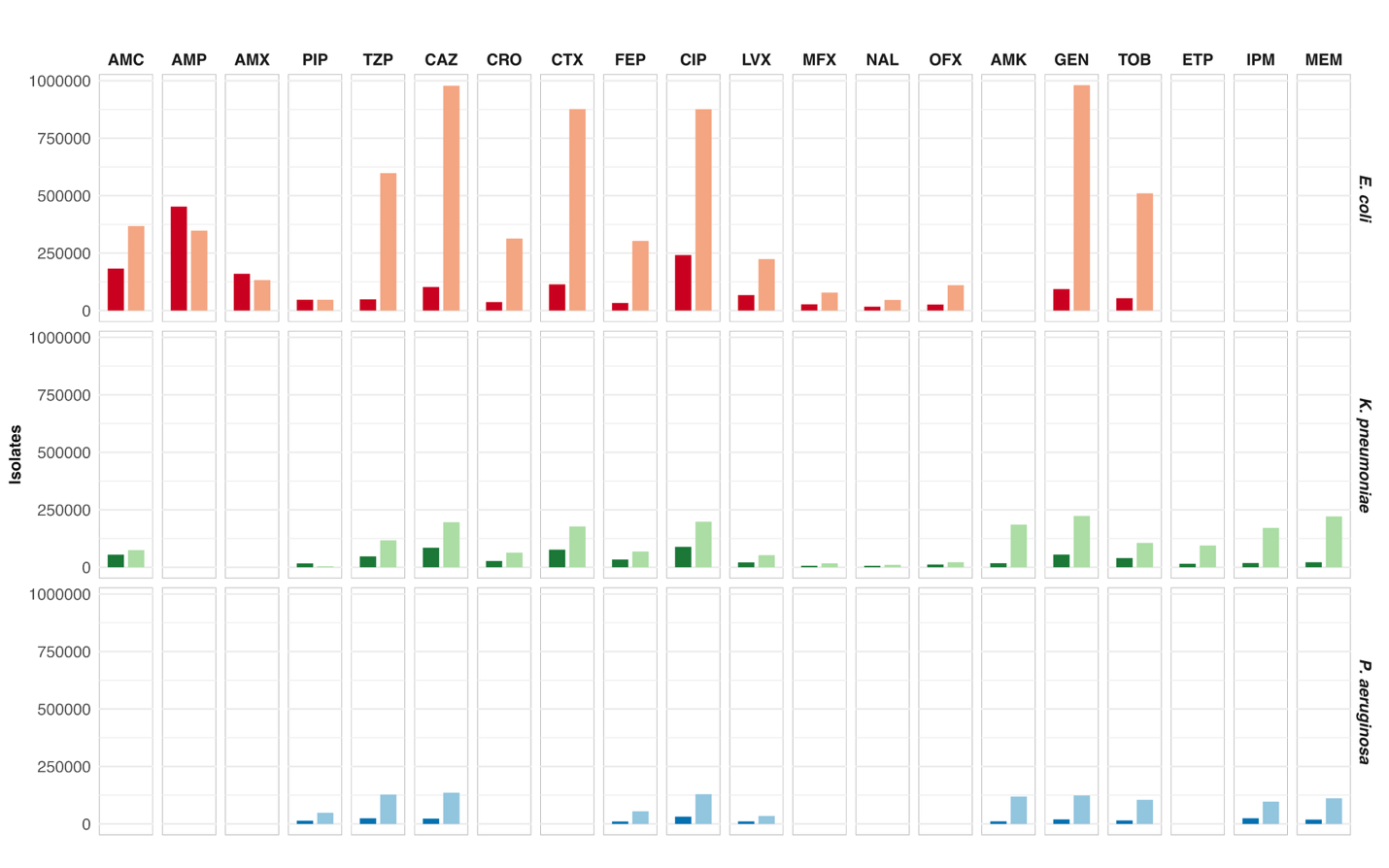

**Figure S1.** The number of susceptible (opaque) and resistant (transparent) bacterial isolates tested for each antibiotic and pathogen in the dataset.

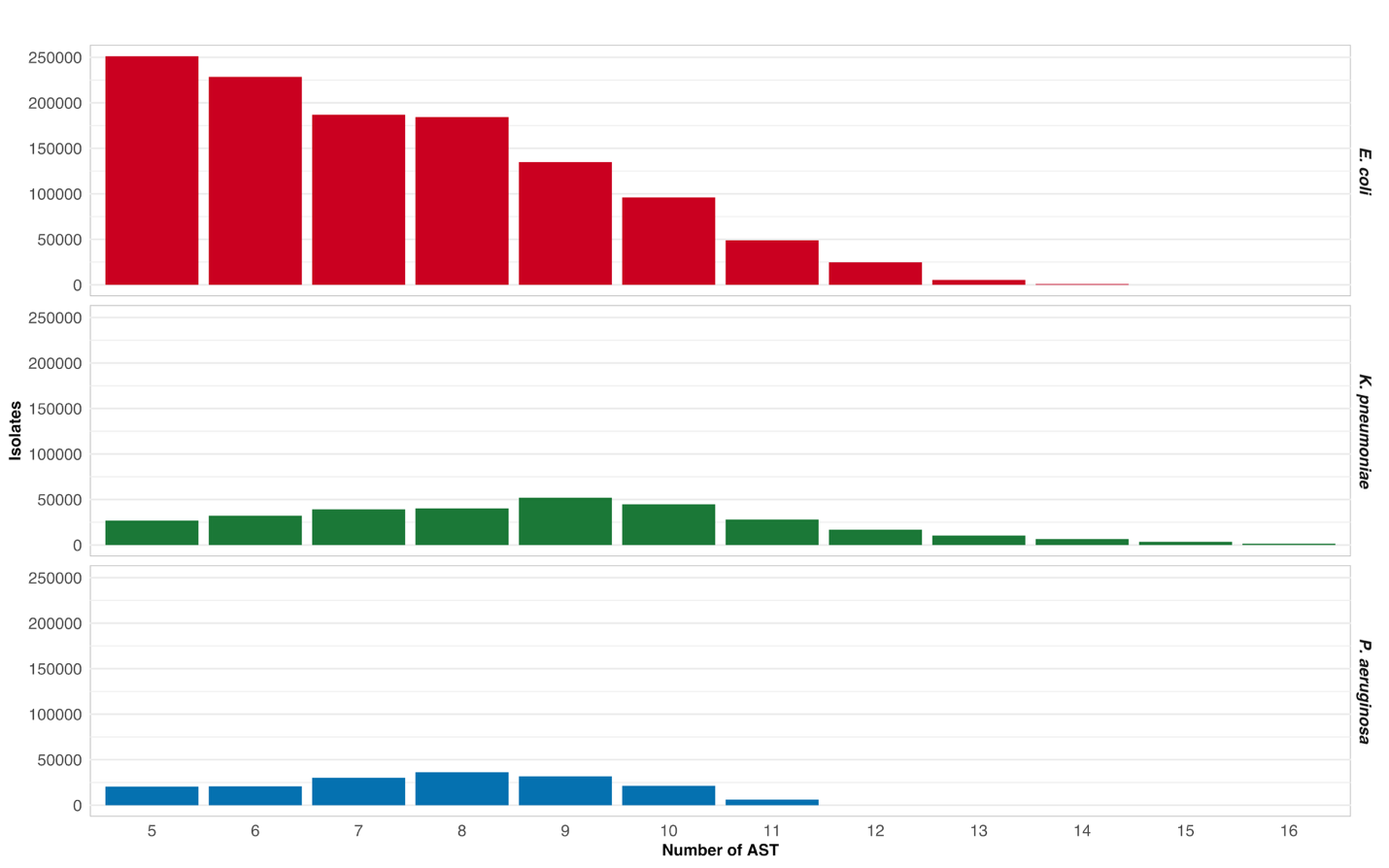

**Figure S2.** Distribution of the number of antibiotics tested per bacterial isolate for each pathogen in the dataset.

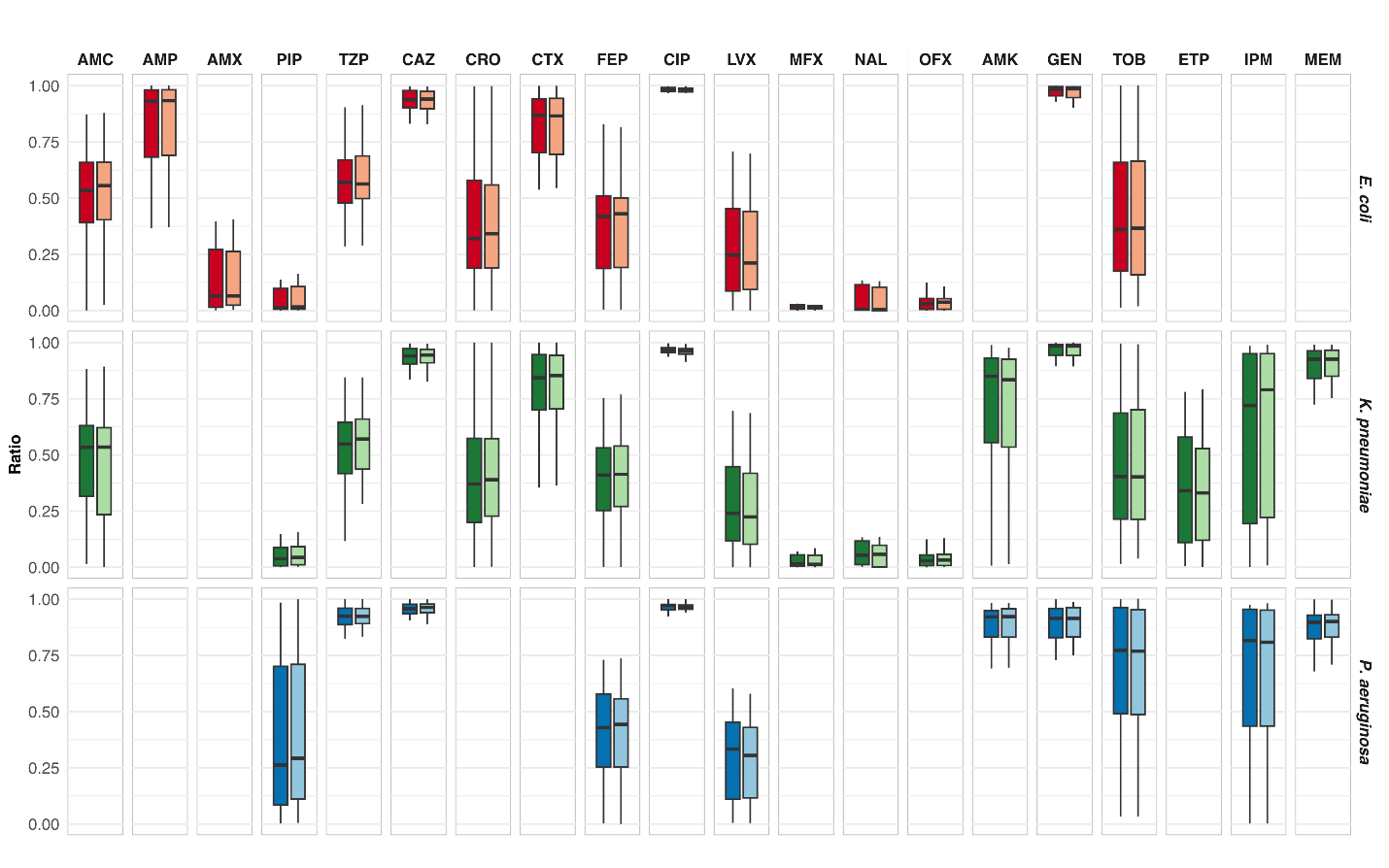

**Figure S3.** Distribution of the proportion of bacterial isolates tested against each antibiotic for female (opaque) and male (transparent) patients and for each pathogen across countries in the dataset. The center line, box limits, and whiskers represent the median, upper, and lower quartiles, and 1.5 times the interquartile range, respectively.

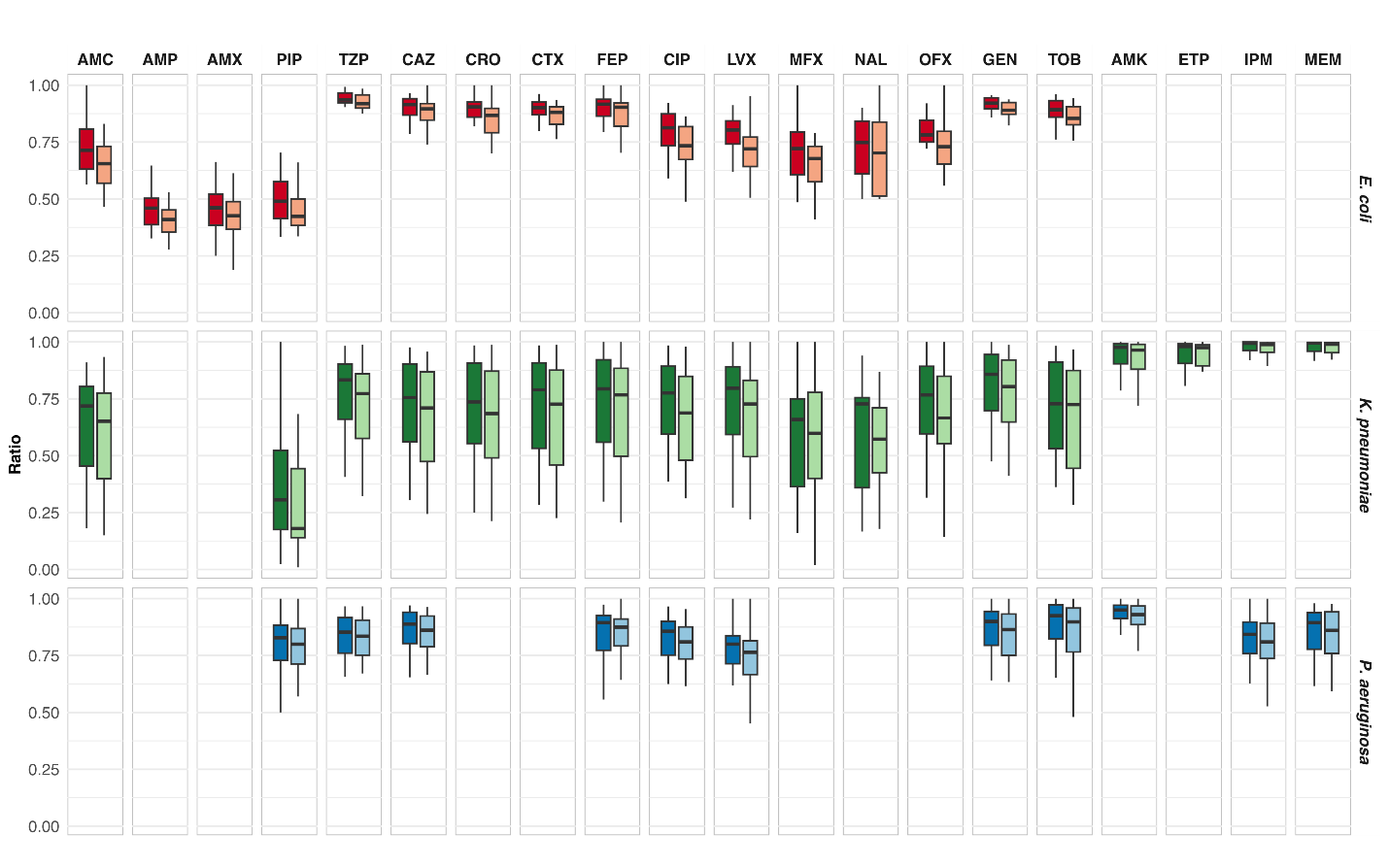

**Figure S4.** The proportion of bacterial isolates susceptible to each antibiotic for female (opaque) and male (transparent) patients and for each pathogen across countries in the dataset. The center line, box limits, and whiskers represent the median, upper, and lower quartiles, and 1.5 times the interquartile range, respectively.

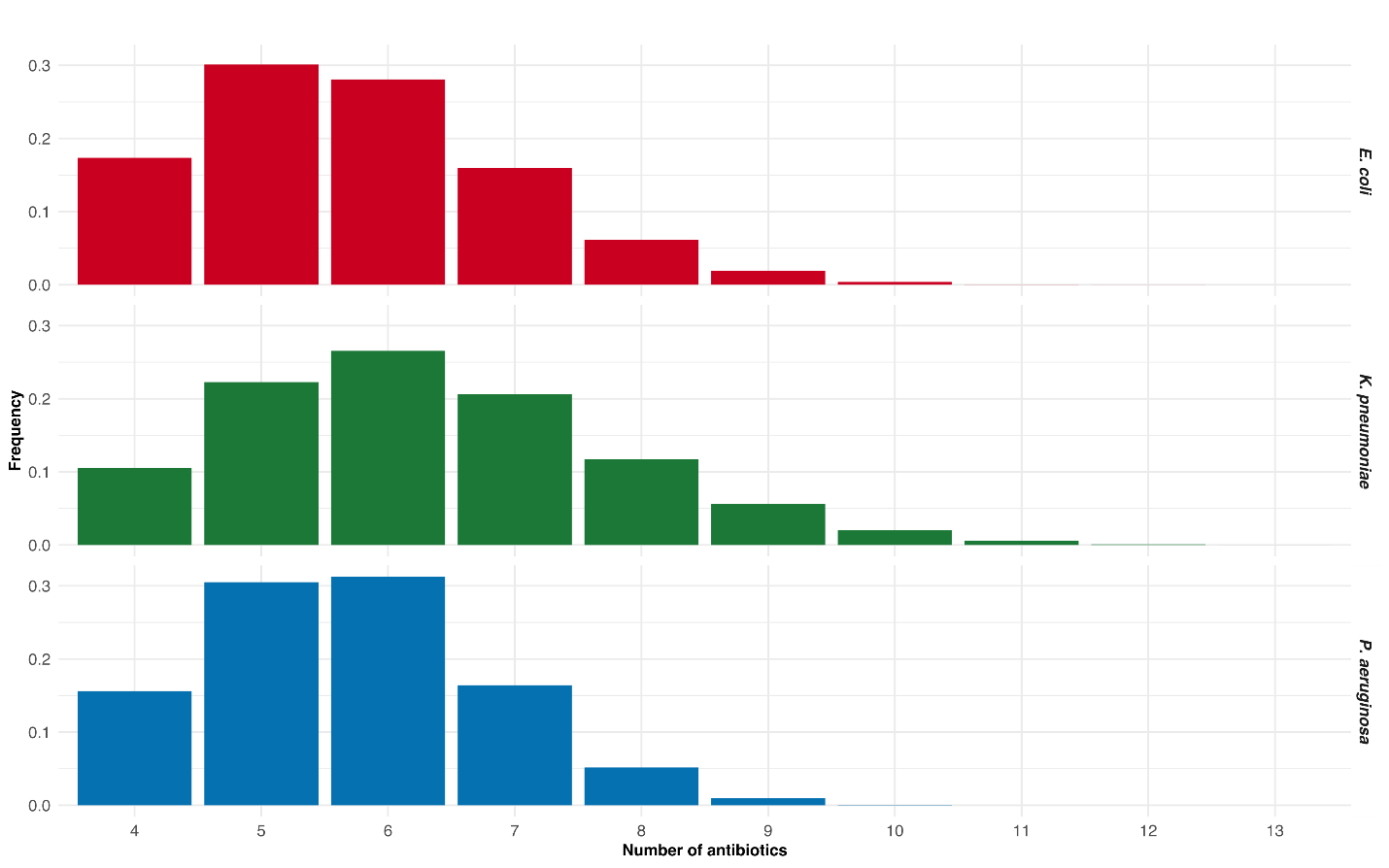

**Figure S5.** Histogram of the number of antibiotic susceptibility testing (AST) results used as input for the model during training at each epoch for each pathogen.

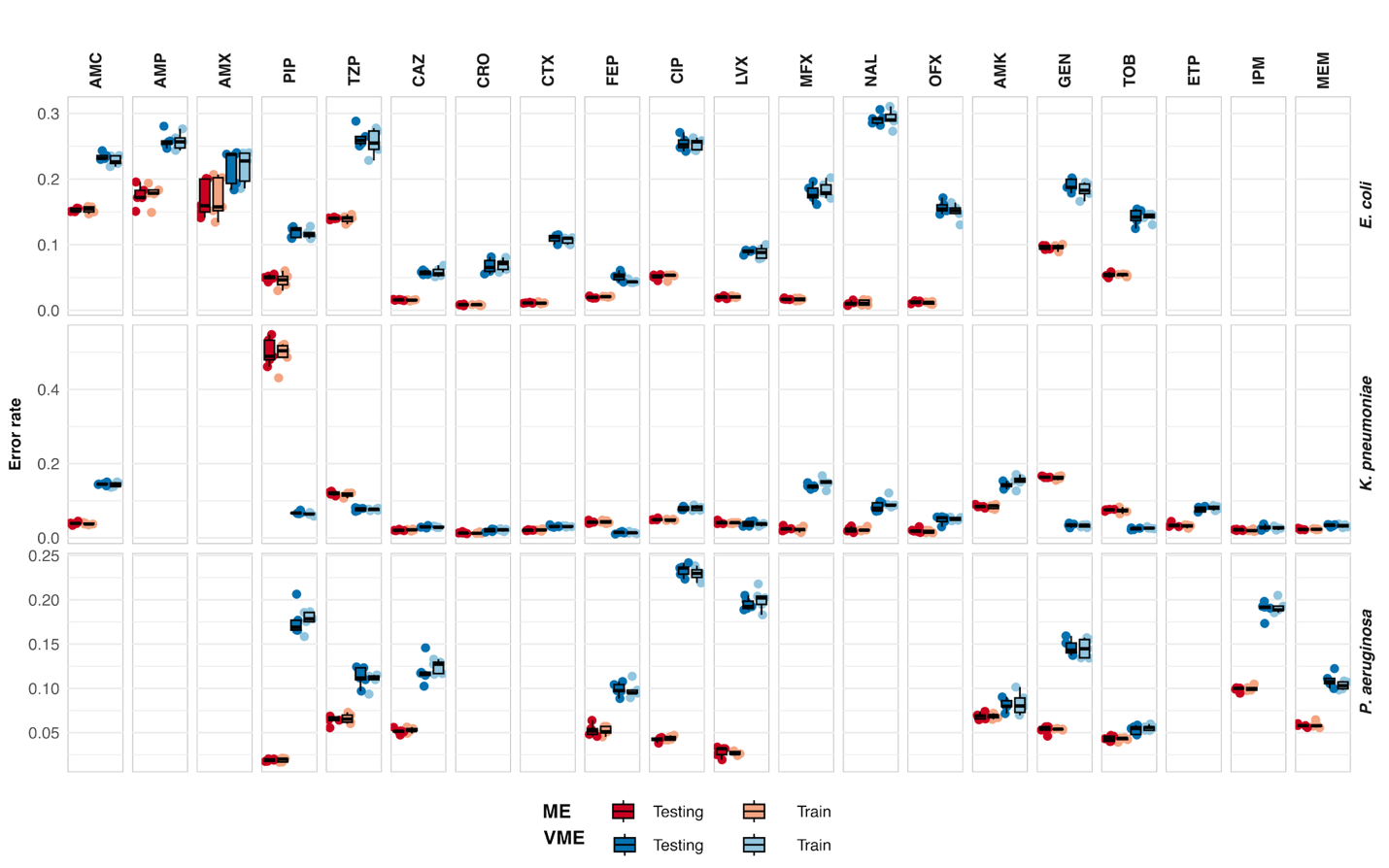

**Figure S6.** Performance of the cross-validation. The major error (red) and very major error (VME) rates for each fold are represented by dots, the horizontal line, box limits, and whiskers represent the median, upper, and lower quartiles, and 1.5 times the interquartile range, respectively, across folds.

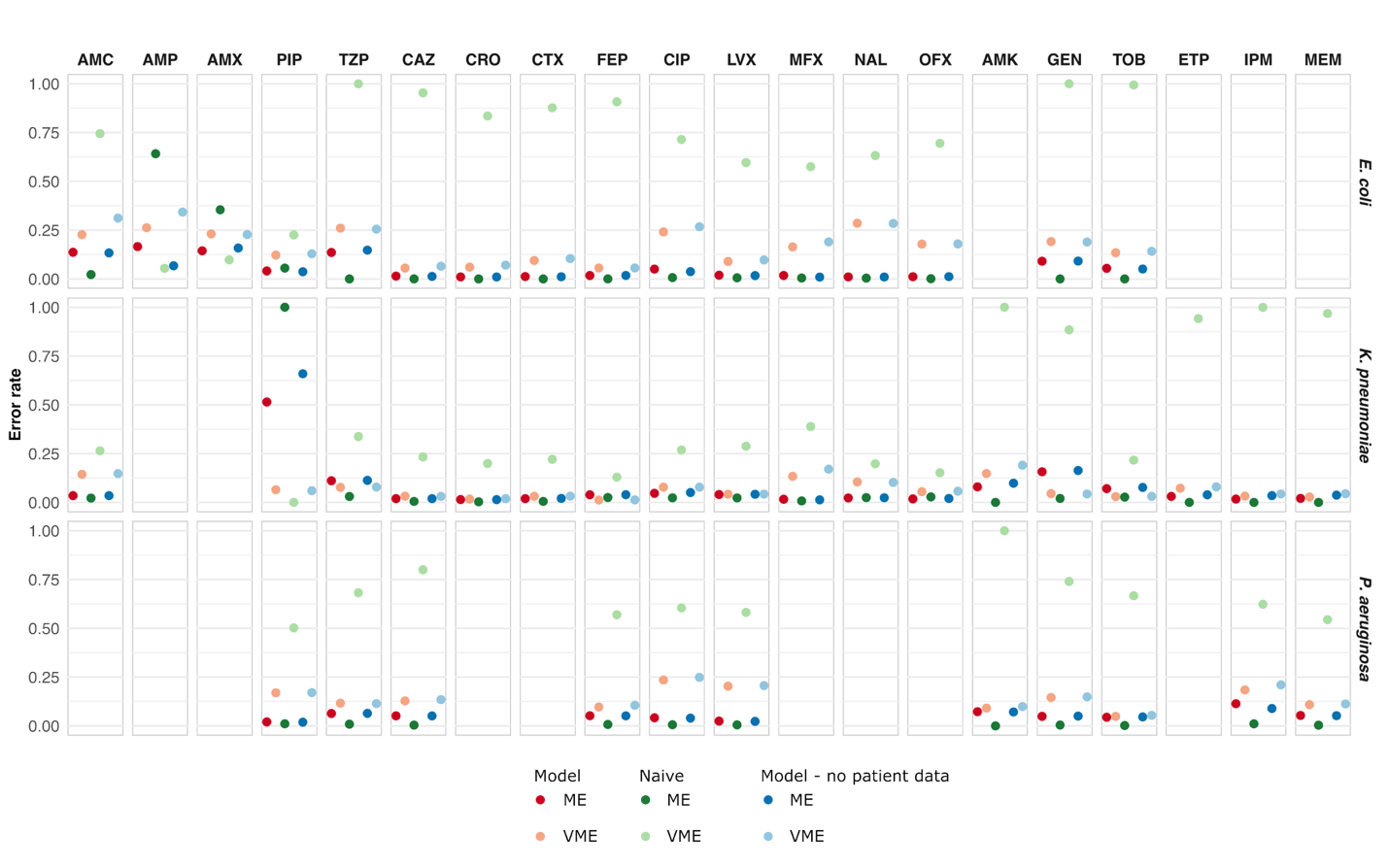

**Figure S7.** Performance of the model with and without patient data and the naive classifier.

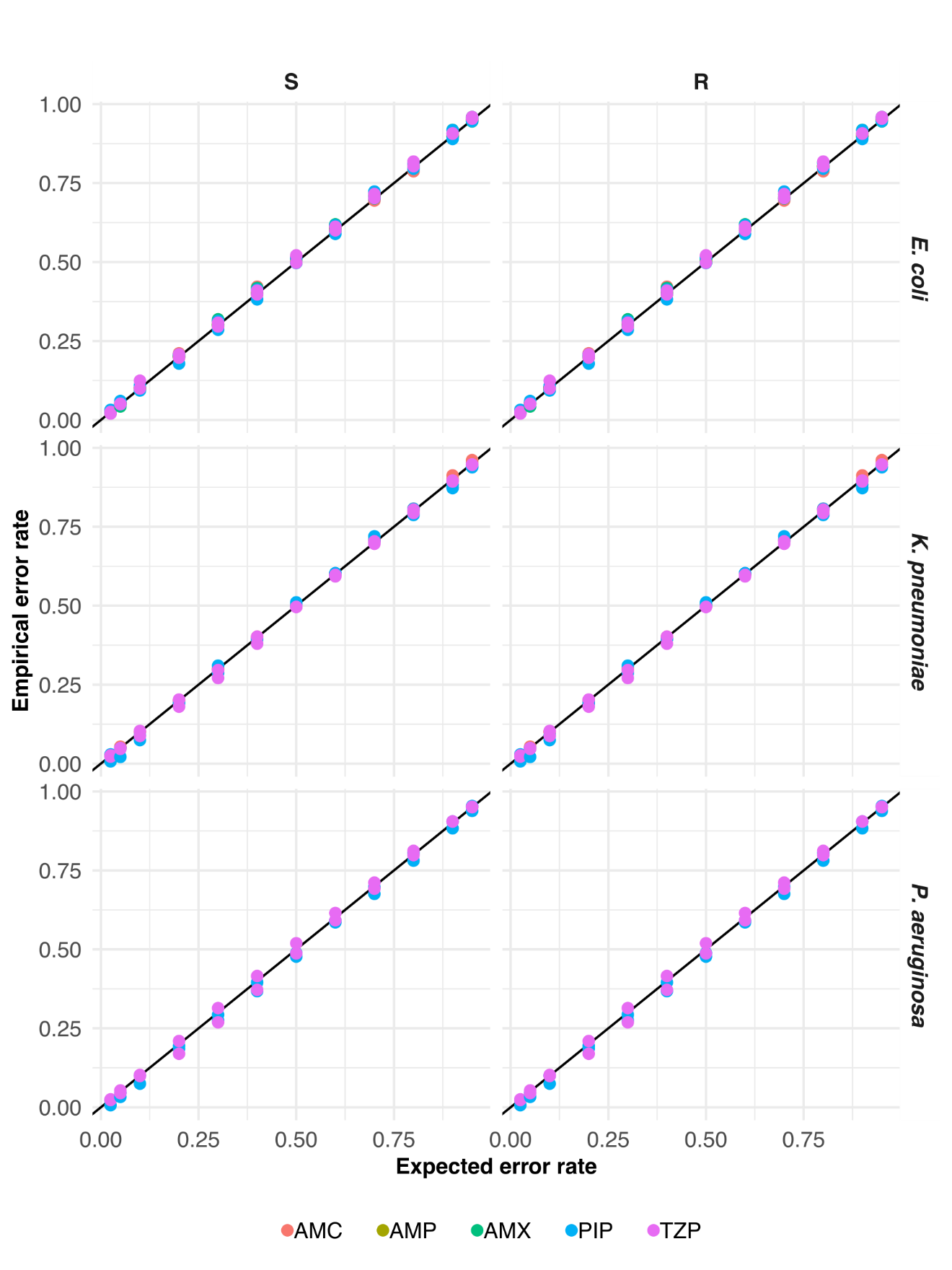

**Figure S8.** Empirical and expected error rates and performance of the model after conditional inductive conformal prediction. For susceptible (S) and resistant (R) pathogens and penicillin antibiotics.

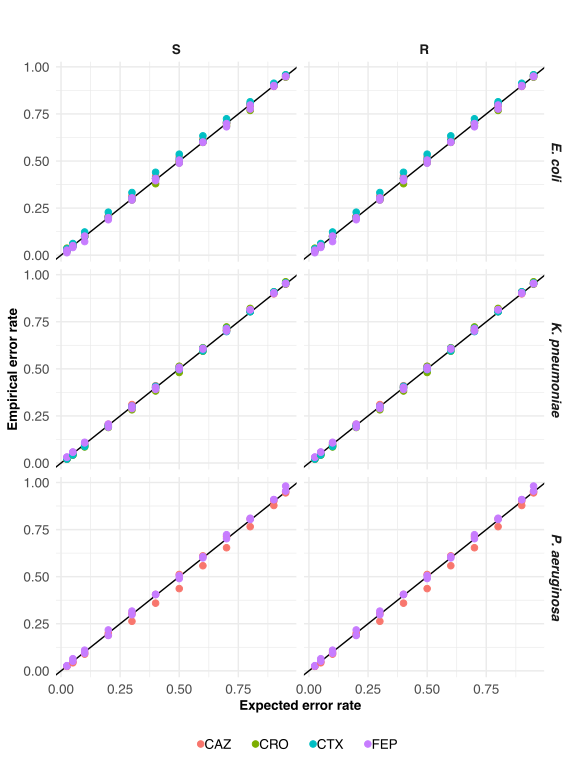

**Figure S9.** Empirical and expected error rates and performance of the model after conditional inductive conformal prediction. For susceptible (S) and resistant (R) pathogens and antibiotics belonging to cephalosporins.

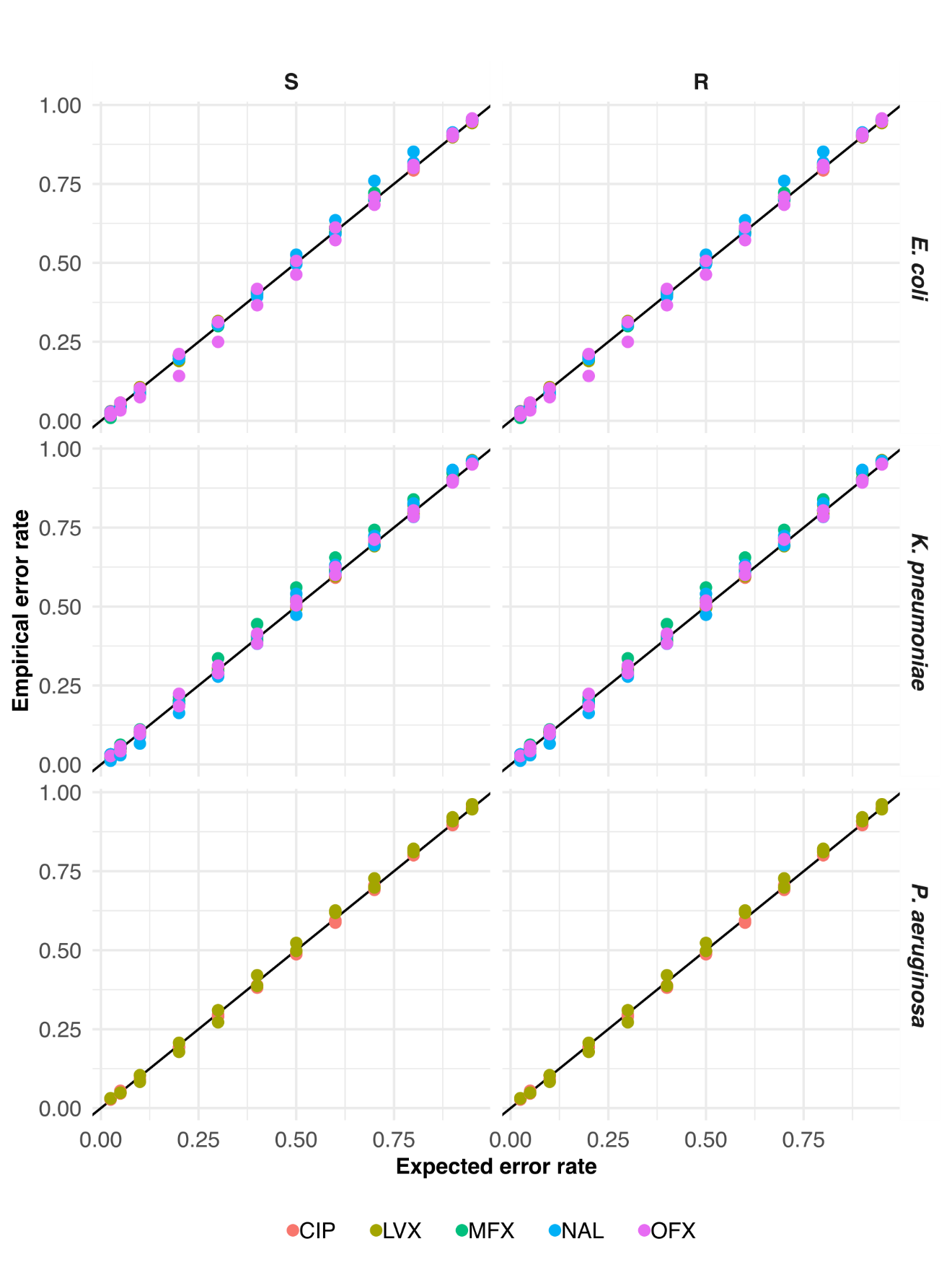

**Figure S10.** Empirical and expected error rates and performance of the model after conditional inductive conformal prediction. For susceptible (S) and resistant (R) pathogens and quinolone antibiotics.

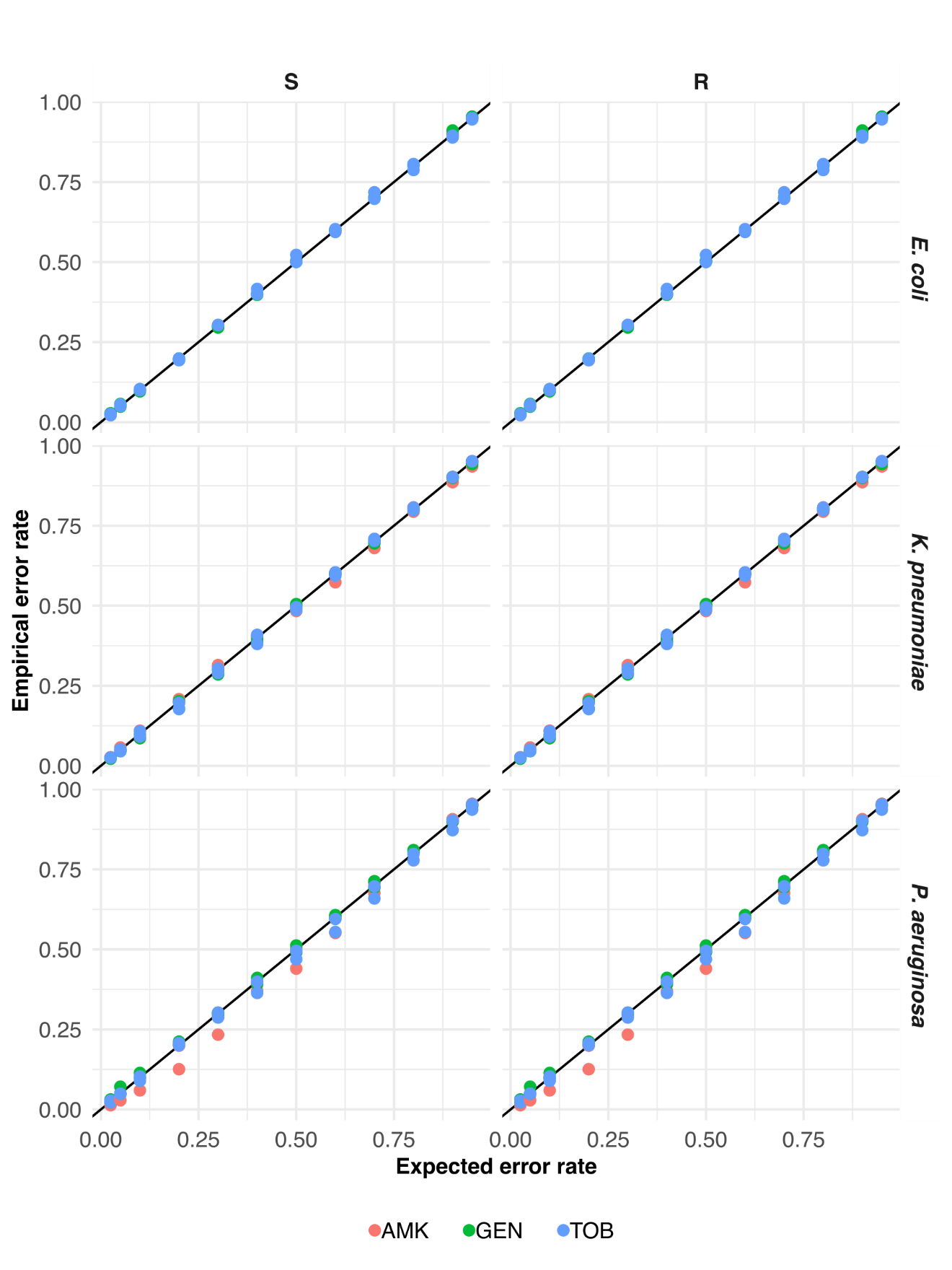

**Figure S11.** Empirical and expected error rates and performance of the model after conditional inductive conformal prediction. For susceptible (S) and resistant (R) pathogens and aminoglycoside antibiotics.

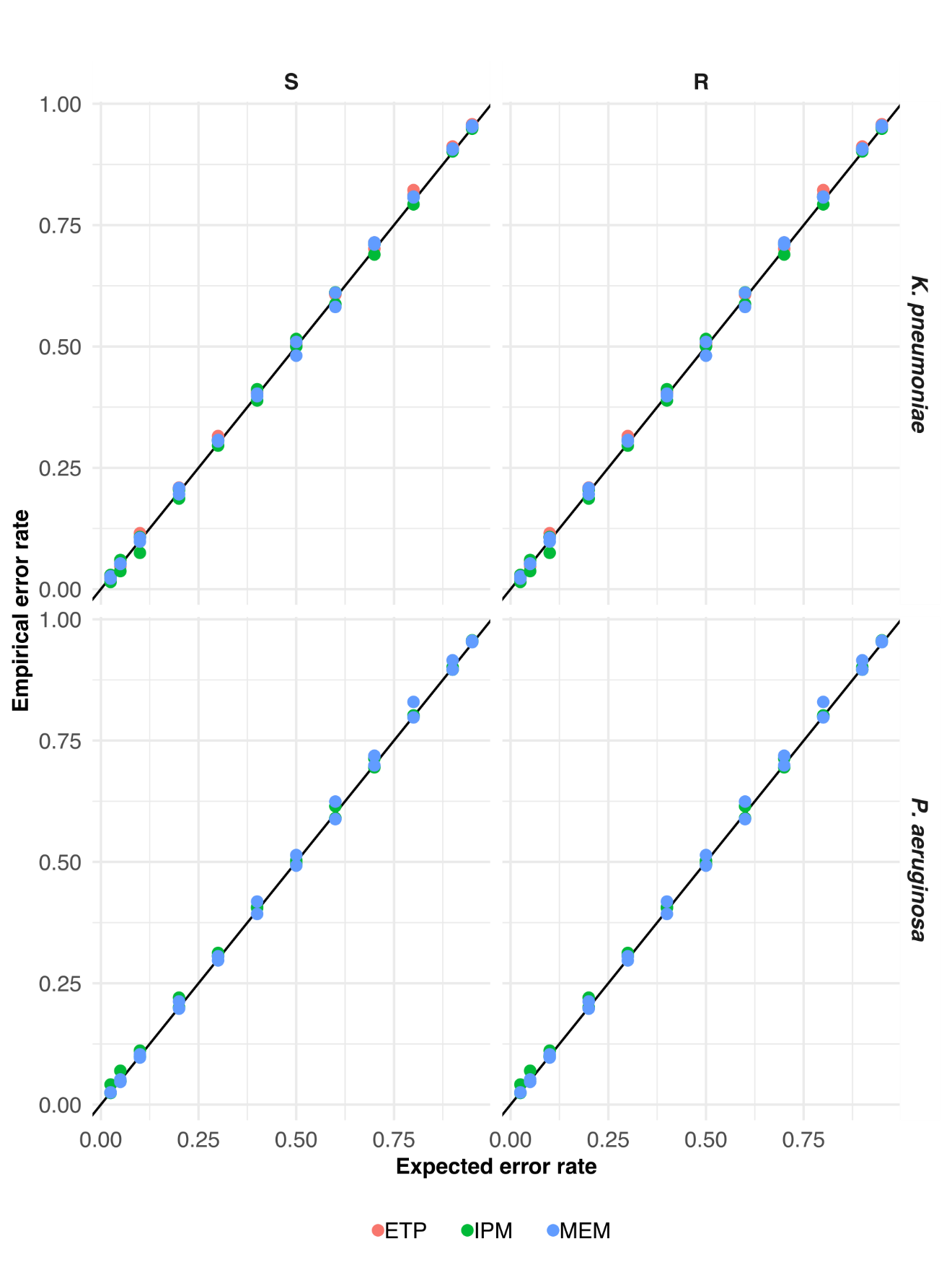

**Figure S12.** Empirical and expected error rates and performance of the model after conditional inductive conformal prediction. For susceptible (S) and resistant (R) pathogens and carbapenem antibiotics.

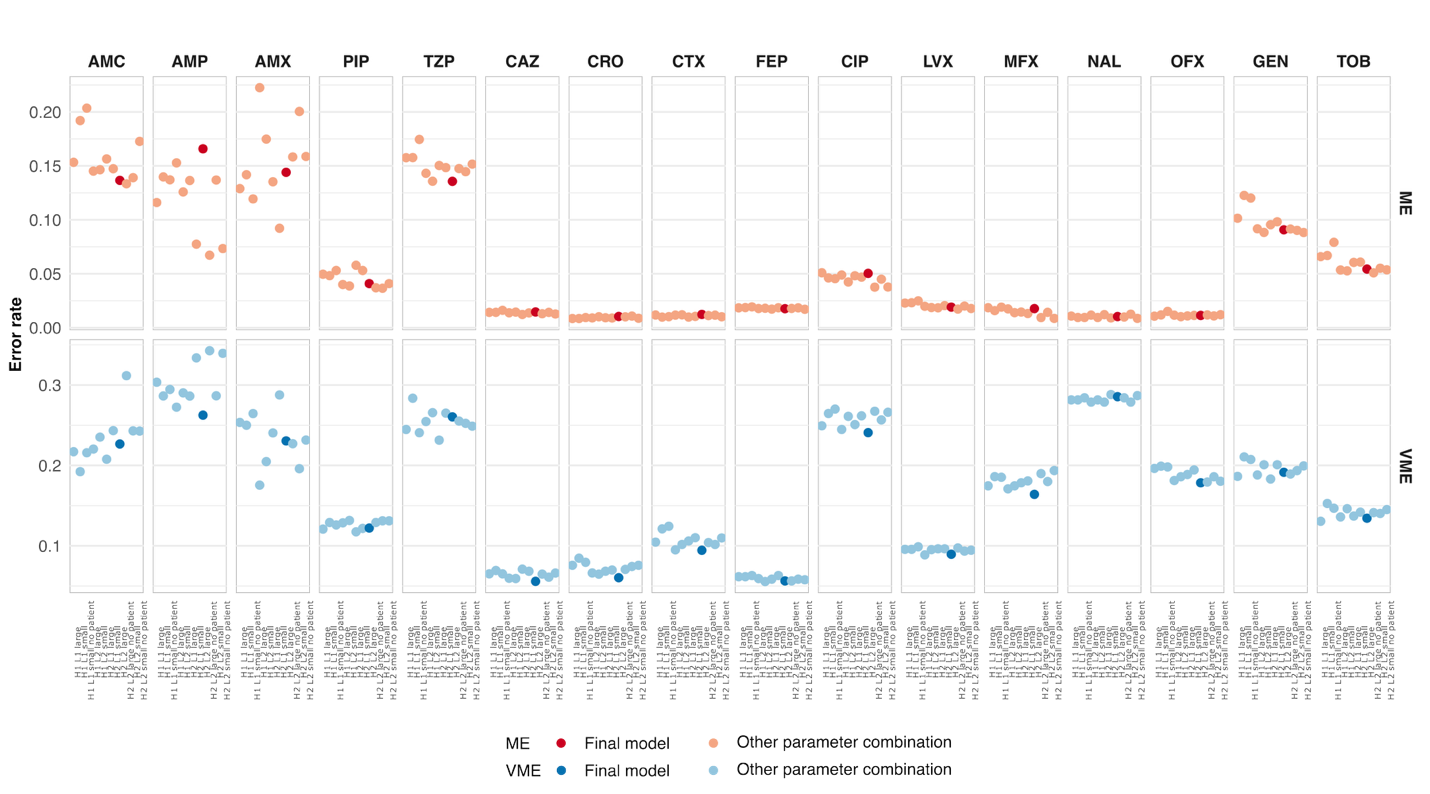

**Figure S13.** Performance of the model with different hyperparameters: 1 or 2 heads (1H, 2H), 1 or 2 layers (1L, 2L), word embedding of size 64 and position-wise feed-forward layer of 128 (small), or word embedding of size 128 and position-wise feed-forward layer of 256 (large), with patient and AST data, or without patient data (no patient), for *E. coli*.

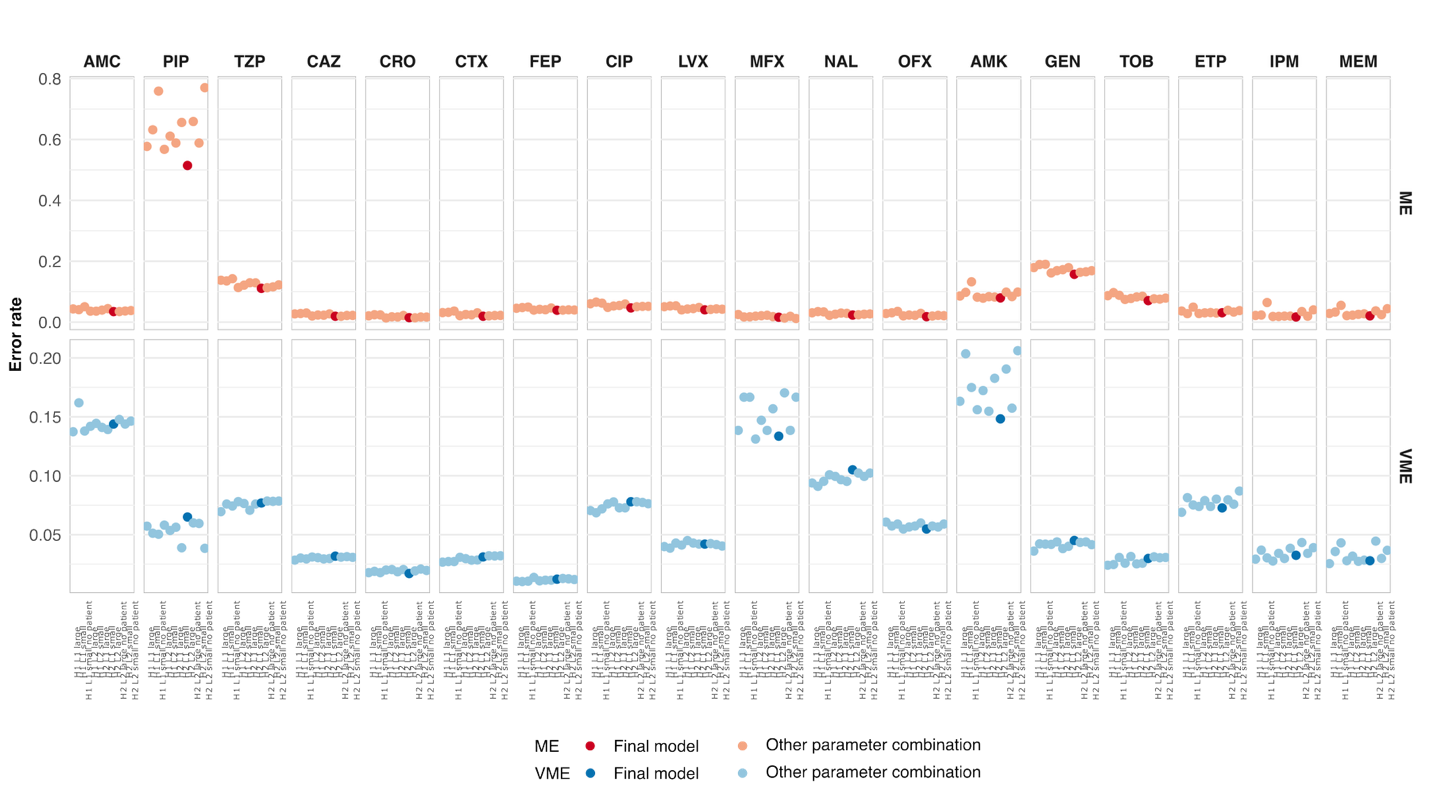

**Figure S14.** Performance of the model with different hyperparameters: 1 or 2 heads (1H, 2H), 1 or 2 layers (1L, 2L), word embedding of size 64 and position-wise feed-forward layer of 128 (small), or word embedding of size 128 and position-wise feed-forward layer of 256 (large), with patient and AST data, or without patient data (no patient), for *K. pneumoniae*, and (S15) *P. aeruginosa*.

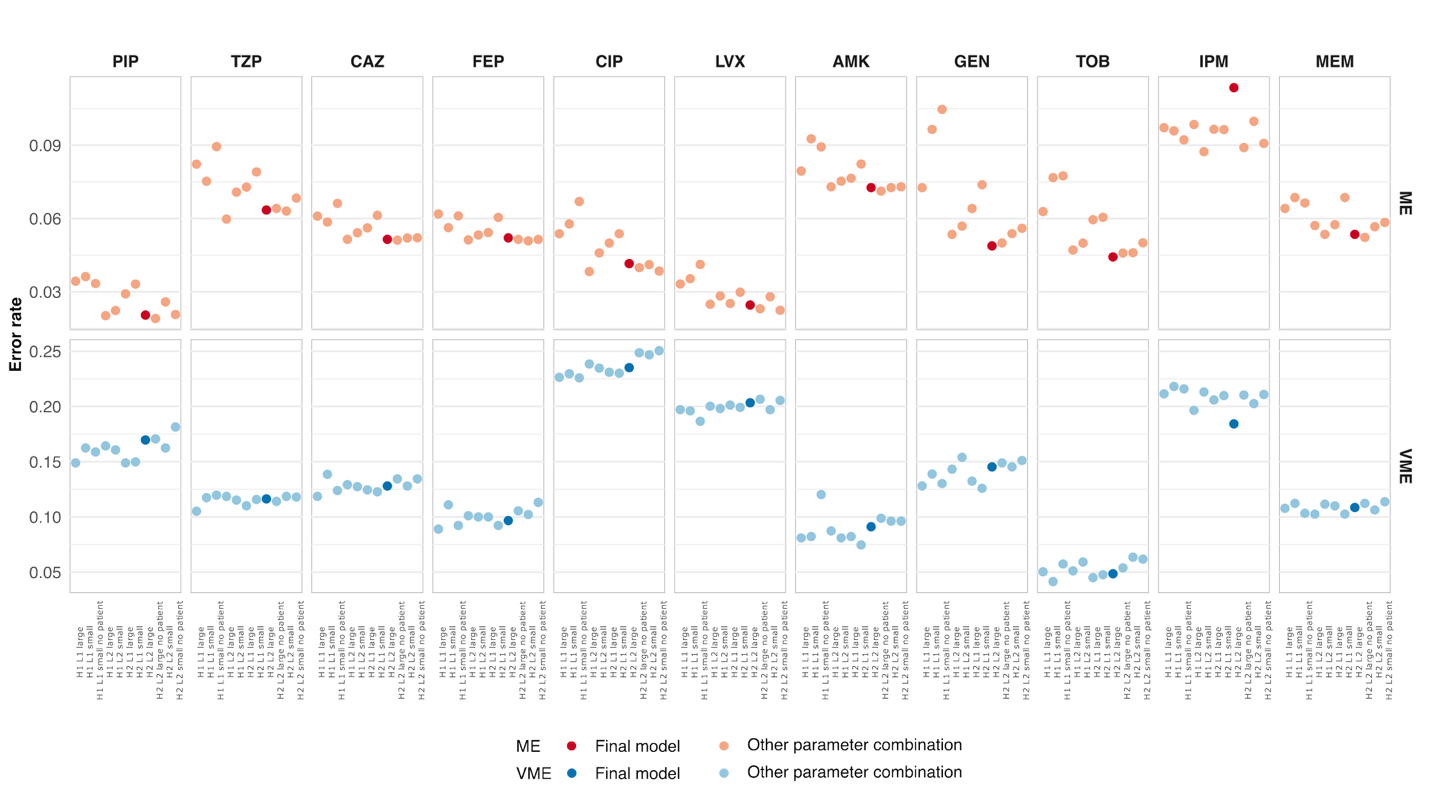

**Figure S15.** Performance of the model with different hyperparameters: 1 or 2 heads (1H, 2H), 1 or 2 layers (1L, 2L), word embedding of size 64 and position-wise feed-forward layer of 128 (small), or word embedding of size 128 and position-wise feed-forward layer of 256 (large), with patient and AST data, or without patient data (no patient), for *P. aeruginosa*.

**Table S1.** **Data expansion.** For isolate *j* with $A_{j}$ AST results, we created a fixed number of data points at each epoch of training, calibration, and testing. The number of AST results, $a_{k}$, included in the input sentence $x_{k}$ of data point $k$ was sampled from a binomial distribution with parameters ${N=A}_{j}-1$ and $p=\frac{D_{j}}{A_{j}-1}-1$, listed below. If $a_{k}<4$, the data point was dismissed. The number of antibiotic susceptibilities to predict for data point $k$ was given by $A_{j}-a_{k}$, and the true susceptibility test results are summarized in the sequence $y_{k}$. The decision on which antibiotics were included in $x_{k}$ and $y_{k}$ was done randomly for all data points.

| **Number of AST results available for isolate *j*,** $\boldsymbol{A}_{\boldsymbol{j}}$ | **Number of data points created per isolate at each epoch** | **Expected number of input AST results per data point,** $\boldsymbol{D}_{\boldsymbol{j}}$ |
| --- | --- | --- |
| 5 | 2 | 4 |
| 6 | 2 | 4.5 |
| 7 | 3 | 5.5 |
| 8 | 3 | 6 |
| 9 | 3 | 6.5 |
| 10 | 4 | 7 |
| 11 | 4 | 8 |
| 12 | 4 | 8 |
| 13 | 4 | 9 |
| 14 | 4 | 9 |
| 15 | 5 | 9 |

**Table S2.** **Performance of the transformer model.** Results from the test dataset: F1-score, major error (ME) rate, and very major error (VME) rate shown for each antibiotic and pathogen, considering all possible numbers of input AST results. Dash (–) represents combinations of bacterial species and antibiotics that the model does not cover.

| Antibiotic | *Escherichia coli* | | | *Klebsiella pneumoniae* | | | *Pseudomonas aeruginosa* | | |
| --- | --- | --- | --- | --- | --- | --- | --- | --- | --- |
|  | F1-score | ME | VME | F1-score | ME | VME | F1-score | ME | VME |
| AMC – Amoxicillin / clavulanic acid | 0.75 | 0.14 | 0.23 | 0.9 | 0.03 | 0.14 | – | – | – |
| AMP – Ampicillin | 0.79 | 0.17 | 0.26 | – | – | – | – | – | – |
| AMX – Amoxicillin | 0.81 | 0.14 | 0.23 | – | – | – | – | – | – |
| PIP – Piperacillin | 0.92 | 0.04 | 0.12 | 0.9 | 0.51 | 0.06 | 0.87 | 0.02 | 0.17 |
| TZP – Piperacillin/ tazobactam | 0.45 | 0.14 | 0.26 | 0.84 | 0.11 | 0.08 | 0.8 | 0.06 | 0.12 |
| CAZ – Ceftazidime | 0.91 | 0.01 | 0.06 | 0.96 | 0.02 | 0.03 | 0.81 | 0.05 | 0.13 |
| CRO – Ceftriaxone | 0.93 | 0.01 | 0.06 | 0.98 | 0.01 | 0.02 | – | – | – |
| CTX – Cefotaxime | 0.9 | 0.01 | 0.09 | 0.96 | 0.02 | 0.03 | – | – | – |
| FEP – Cefepime | 0.89 | 0.02 | 0.06 | 0.96 | 0.04 | 0.01 | 0.83 | 0.05 | 0.1 |
| CIP – Ciprofloxacin | 0.78 | 0.05 | 0.24 | 0.91 | 0.05 | 0.08 | 0.79 | 0.04 | 0.24 |
| LVX – Levofloxacin | 0.92 | 0.02 | 0.09 | 0.93 | 0.04 | 0.04 | 0.85 | 0.02 | 0.2 |
| MFX – Moxifloxacin | 0.89 | 0.02 | 0.16 | 0.91 | 0.02 | 0.13 | – | – | – |
| NAL – Nalidixic acid | 0.82 | 0.01 | 0.29 | 0.93 | 0.02 | 0.11 | – | – | – |
| OFX – Ofloxacin | 0.88 | 0.01 | 0.18 | 0.96 | 0.02 | 0.05 | – | – | – |
| AMK – Amikacin | – | – | – | 0.65 | 0.08 | 0.15 | 0.66 | 0.07 | 0.09 |
| GEN – Gentamicin | 0.58 | 0.09 | 0.19 | 0.74 | 0.16 | 0.05 | 0.79 | 0.05 | 0.15 |
| TOB – Tobramycin | 0.73 | 0.05 | 0.13 | 0.9 | 0.07 | 0.03 | 0.84 | 0.04 | 0.05 |
| ETP – Ertapenem | – | – | – | 0.88 | 0.03 | 0.07 | – | – | – |
| IPM – Imipenem | – | – | – | 0.92 | 0.02 | 0.03 | 0.71 | 0.11 | 0.18 |
| MEM – Meropenem | – | – | – | 0.91 | 0.02 | 0.03 | 0.81 | 0.05 | 0.11 |

**Table S3.** **Performance of the transformer model as a function of the number of antibiotic susceptibility testing (AST) results included in the input.** Results from the test dataset: F1-score, major error (ME) rate, and very major error (VME) rate shown for each antibiotic, pathogen, and the number of antibiotic susceptibility testing (AST) results included in the input. Dash (–) indicates combinations of bacterial species and antibiotics that the model does not cover. We limited the table to 8 AST results in the input due to the low number of isolates tested for 10 or more antibiotics in the data.

| Antibiotic | AST results in input | *Escherichia coli* | | | *Klebsiella pneumoniae* | | | *Pseudomonas aeruginosa* | | |
| --- | --- | --- | --- | --- | --- | --- | --- | --- | --- | --- |
|  |  | F1 score | ME | VME | F1 score | ME | VME | F1 score | ME | VME |
| AMC – Amoxicillin / clavulanic acid | 4 | 0.71 | 0.15 | 0.29 | 0.89 | 0.04 | 0.15 | – | – | – |
|  | 5 | 0.74 | 0.14 | 0.26 | 0.89 | 0.03 | 0.17 | – | – | – |
|  | 6 | 0.75 | 0.13 | 0.22 | 0.89 | 0.04 | 0.15 | – | – | – |
|  | 7 | 0.77 | 0.13 | 0.19 | 0.9 | 0.03 | 0.14 | – | – | – |
|  | 8 | 0.8 | 0.13 | 0.14 | 0.91 | 0.03 | 0.13 | – | – | – |
| AMP – Ampicillin | 4 | 0.7 | 0.22 | 0.37 | – | – | – | – | – | – |
|  | 5 | 0.78 | 0.23 | 0.24 | – | – | – | – | – | – |
|  | 6 | 0.81 | 0.13 | 0.26 | – | – | – | – | – | – |
|  | 7 | 0.85 | 0.07 | 0.21 | – | – | – | – | – | – |
|  | 8 | 0.89 | 0.07 | 0.15 | – | – | – | – | – | – |
| AMX – Amoxicillin | 4 | 0.74 | 0.34 | 0.22 | – | – | – | – | – | – |
|  | 5 | 0.78 | 0.16 | 0.27 | – | – | – | – | – | – |
|  | 6 | 0.82 | 0.11 | 0.24 | – | – | – | – | – | – |
|  | 7 | 0.86 | 0.07 | 0.19 | – | – | – | – | – | – |
|  | 8 | 0.89 | 0.06 | 0.14 | – | – | – | – | – | – |
| PIP – Piperacillin | 4 | 0.81 | 0.09 | 0.25 | 0.86 | 0.74 | 0.05 | 0.83 | 0.03 | 0.2 |
|  | 5 | 0.9 | 0.03 | 0.17 | 0.89 | 0.7 | 0.07 | 0.87 | 0.03 | 0.17 |
|  | 6 | 0.92 | 0.05 | 0.12 | 0.91 | 0.59 | 0.04 | 0.9 | 0.01 | 0.14 |
|  | 7 | 0.93 | 0.04 | 0.09 | 0.9 | 0.53 | 0.05 | 0.86 | 0.02 | 0.18 |
|  | 8 | 0.97 | 0.02 | 0.03 | 0.91 | 0.39 | 0.09 | 0.88 | 0.01 | 0.19 |
| TZP – Piperacillin/ tazobactam | 4 | 0.43 | 0.12 | 0.31 | 0.82 | 0.11 | 0.11 | 0.76 | 0.05 | 0.19 |
|  | 5 | 0.43 | 0.14 | 0.29 | 0.82 | 0.11 | 0.09 | 0.78 | 0.07 | 0.13 |
|  | 6 | 0.44 | 0.14 | 0.25 | 0.83 | 0.12 | 0.09 | 0.82 | 0.06 | 0.09 |
|  | 7 | 0.47 | 0.13 | 0.2 | 0.85 | 0.12 | 0.06 | 0.81 | 0.07 | 0.1 |
|  | 8 | 0.54 | 0.11 | 0.2 | 0.86 | 0.1 | 0.07 | 0.88 | 0.04 | 0.01 |
| CAZ – Ceftazidime | 4 | 0.88 | 0.02 | 0.08 | 0.93 | 0.03 | 0.07 | 0.78 | 0.04 | 0.2 |
|  | 5 | 0.9 | 0.02 | 0.06 | 0.96 | 0.02 | 0.04 | 0.8 | 0.05 | 0.14 |
|  | 6 | 0.91 | 0.01 | 0.05 | 0.96 | 0.02 | 0.03 | 0.82 | 0.05 | 0.1 |
|  | 7 | 0.93 | 0.01 | 0.03 | 0.97 | 0.02 | 0.02 | 0.79 | 0.06 | 0.11 |
|  | 8 | 0.96 | 0.01 | 0.01 | 0.98 | 0.01 | 0.01 | 0.86 | 0.04 | 0.04 |
| CRO – Ceftriaxone | 4 | 0.89 | 0.02 | 0.08 | 0.95 | 0.02 | 0.05 | – | – | – |
|  | 5 | 0.9 | 0.02 | 0.08 | 0.96 | 0.02 | 0.03 | – | – | – |
|  | 6 | 0.93 | 0.01 | 0.06 | 0.98 | 0.02 | 0.01 | – | – | – |
|  | 7 | 0.97 | 0 | 0.04 | 0.99 | 0.01 | 0.01 | – | – | – |
|  | 8 | 0.98 | 0 | 0.02 | 0.98 | 0.01 | 0.02 | – | – | – |
| CTX – Cefotaxime | 4 | 0.86 | 0.02 | 0.13 | 0.93 | 0.03 | 0.08 | – | – | – |
|  | 5 | 0.89 | 0.02 | 0.1 | 0.95 | 0.03 | 0.04 | – | – | – |
|  | 6 | 0.92 | 0.01 | 0.09 | 0.97 | 0.02 | 0.02 | – | – | – |
|  | 7 | 0.96 | 0 | 0.05 | 0.97 | 0.01 | 0.02 | – | – | – |
|  | 8 | 0.97 | 0 | 0.02 | 0.98 | 0.01 | 0.02 | – | – | – |
| FEP – Cefepime | 4 | 0.86 | 0.02 | 0.09 | 0.94 | 0.05 | 0.02 | 0.82 | 0.06 | 0.1 |
|  | 5 | 0.87 | 0.02 | 0.07 | 0.95 | 0.04 | 0.01 | 0.81 | 0.05 | 0.12 |
|  | 6 | 0.9 | 0.02 | 0.05 | 0.95 | 0.04 | 0.01 | 0.85 | 0.05 | 0.08 |
|  | 7 | 0.93 | 0.01 | 0.03 | 0.96 | 0.03 | 0.01 | 0.8 | 0.06 | 0.1 |
|  | 8 | 0.9 | 0.02 | 0.04 | 0.95 | 0.04 | 0.01 | 0.84 | 0.04 | 0.09 |
| CIP – Ciprofloxacin | 4 | 0.69 | 0.05 | 0.36 | 0.86 | 0.06 | 0.13 | 0.73 | 0.04 | 0.32 |
|  | 5 | 0.76 | 0.06 | 0.26 | 0.89 | 0.05 | 0.1 | 0.77 | 0.05 | 0.26 |
|  | 6 | 0.8 | 0.05 | 0.22 | 0.91 | 0.05 | 0.08 | 0.79 | 0.04 | 0.22 |
|  | 7 | 0.85 | 0.04 | 0.15 | 0.93 | 0.04 | 0.06 | 0.85 | 0.04 | 0.14 |
|  | 8 | 0.92 | 0.02 | 0.07 | 0.95 | 0.03 | 0.04 | 0.88 | 0.03 | 0.09 |
| LVX – Levofloxacin | 4 | 0.84 | 0.03 | 0.19 | 0.9 | 0.04 | 0.08 | 0.79 | 0.04 | 0.27 |
|  | 5 | 0.89 | 0.03 | 0.12 | 0.91 | 0.06 | 0.06 | 0.85 | 0.02 | 0.2 |
|  | 6 | 0.94 | 0.02 | 0.08 | 0.92 | 0.05 | 0.05 | 0.83 | 0.03 | 0.22 |
|  | 7 | 0.96 | 0.01 | 0.05 | 0.95 | 0.03 | 0.04 | 0.87 | 0.02 | 0.18 |
|  | 8 | 0.96 | 0.01 | 0.05 | 0.97 | 0.02 | 0.02 | 0.89 | 0.01 | 0.14 |
| MFX – Moxifloxacin | 4 | 0.85 | 0.03 | 0.2 | 0.78 | 0.04 | 0.29 | – | – | – |
|  | 5 | 0.84 | 0.02 | 0.22 | 0.89 | 0.02 | 0.16 | – | – | – |
|  | 6 | 0.87 | 0.02 | 0.18 | 0.88 | 0.02 | 0.18 | – | – | – |
|  | 7 | 0.91 | 0.01 | 0.14 | 0.9 | 0 | 0.18 | – | – | – |
|  | 8 | 0.95 | 0 | 0.09 | 0.96 | 0.02 | 0.05 | – | – | – |
| NAL – Nalidixic acid | 4 | 0.76 | 0.01 | 0.37 | 0.87 | 0.02 | 0.2 | – | – | – |
|  | 5 | 0.78 | 0.01 | 0.33 | 0.91 | 0.03 | 0.12 | – | – | – |
|  | 6 | 0.81 | 0.01 | 0.29 | 0.92 | 0.03 | 0.1 | – | – | – |
|  | 7 | 0.85 | 0.01 | 0.24 | 0.93 | 0.03 | 0.1 | – | – | – |
|  | 8 | 0.87 | 0 | 0.23 | 0.92 | 0 | 0.14 | – | – | – |
| OFX – Ofloxacin | 4 | 0.79 | 0.01 | 0.31 | 0.92 | 0.02 | 0.11 | – | – | – |
|  | 5 | 0.83 | 0.01 | 0.26 | 0.94 | 0.02 | 0.08 | – | – | – |
|  | 6 | 0.88 | 0.01 | 0.18 | 0.96 | 0.02 | 0.05 | – | – | – |
|  | 7 | 0.94 | 0.01 | 0.08 | 0.96 | 0.02 | 0.06 | – | – | – |
|  | 8 | 0.93 | 0.02 | 0.06 | 0.99 | 0 | 0.02 | – | – | – |
| AMK – Amikacin | 4 | – | – | – | 0.6 | 0.08 | 0.19 | 0.63 | 0.08 | 0.12 |
|  | 5 | – | – | – | 0.64 | 0.07 | 0.18 | 0.67 | 0.07 | 0.11 |
|  | 6 | – | – | – | 0.6 | 0.08 | 0.19 | 0.66 | 0.07 | 0.09 |
|  | 7 | – | – | – | 0.68 | 0.07 | 0.11 | 0.66 | 0.08 | 0.04 |
|  | 8 | – | – | – | 0.7 | 0.09 | 0.09 | 0.69 | 0.05 | 0.03 |
| GEN – Gentamicin | 4 | 0.5 | 0.12 | 0.24 | 0.71 | 0.16 | 0.07 | 0.76 | 0.05 | 0.15 |
|  | 5 | 0.59 | 0.09 | 0.19 | 0.75 | 0.15 | 0.05 | 0.78 | 0.05 | 0.15 |
|  | 6 | 0.61 | 0.09 | 0.17 | 0.72 | 0.17 | 0.04 | 0.79 | 0.05 | 0.15 |
|  | 7 | 0.6 | 0.07 | 0.21 | 0.73 | 0.16 | 0.04 | 0.81 | 0.04 | 0.13 |
|  | 8 | 0.62 | 0.07 | 0.15 | 0.77 | 0.14 | 0.04 | 0.83 | 0.03 | 0.12 |
| TOB – Tobramycin | 4 | 0.66 | 0.07 | 0.17 | 0.89 | 0.07 | 0.04 | 0.82 | 0.04 | 0.12 |
|  | 5 | 0.72 | 0.06 | 0.14 | 0.9 | 0.06 | 0.04 | 0.82 | 0.05 | 0.05 |
|  | 6 | 0.74 | 0.05 | 0.13 | 0.9 | 0.08 | 0.03 | 0.85 | 0.05 | 0.02 |
|  | 7 | 0.77 | 0.05 | 0.09 | 0.91 | 0.08 | 0.03 | 0.85 | 0.05 | 0.02 |
|  | 8 | 0.77 | 0.05 | 0.1 | 0.91 | 0.08 | 0.02 | 0.86 | 0.03 | 0.08 |
| ETP – Ertapenem | 4 | – | – | – | 0.82 | 0.04 | 0.1 | – | – | – |
|  | 5 | – | – | – | 0.84 | 0.04 | 0.1 | – | – | – |
|  | 6 | – | – | – | 0.86 | 0.04 | 0.08 | – | – | – |
|  | 7 | – | – | – | 0.88 | 0.03 | 0.09 | – | – | – |
|  | 8 | – | – | – | 0.93 | 0.02 | 0.04 | – | – | – |
| IPM – Imipenem | 4 | – | – | – | 0.89 | 0.02 | 0.08 | 0.6 | 0.19 | 0.19 |
|  | 5 | – | – | – | 0.88 | 0.02 | 0.04 | 0.69 | 0.14 | 0.18 |
|  | 6 | – | – | – | 0.91 | 0.02 | 0.04 | 0.75 | 0.09 | 0.17 |
|  | 7 | – | – | – | 0.94 | 0.01 | 0.03 | 0.73 | 0.08 | 0.21 |
|  | 8 | – | – | – | 0.95 | 0.01 | 0.02 | 0.84 | 0.03 | 0.19 |
| MEM – Meropenem | 4 | – | – | – | 0.84 | 0.02 | 0.08 | 0.69 | 0.06 | 0.17 |
|  | 5 | – | – | – | 0.87 | 0.02 | 0.03 | 0.79 | 0.06 | 0.12 |
|  | 6 | – | – | – | 0.9 | 0.02 | 0.02 | 0.83 | 0.05 | 0.09 |
|  | 7 | – | – | – | 0.92 | 0.02 | 0.03 | 0.86 | 0.04 | 0.1 |
|  | 8 | – | – | – | 0.96 | 0.01 | 0.02 | 0.83 | 0.04 | 0.1 |
